# The genetic architecture of leaf vein density traits and its importance for photosynthesis in maize

**DOI:** 10.64898/2026.01.14.699362

**Authors:** José Luis Coyac-Rodríguez, Sergio Pérez-Limón, Elizabeth Hernández-Jaimes, Marcela Hernández-Coronado, Daniel Camo-Escobar, Ana Laura Alonso-Nieves, María de Jesús Ortega-Estrada, Nicole Gómez-Capetillo, Ruairidh James Hay Sawers, Carlos Humberto Ortiz-Ramírez

## Abstract

- Leaf venation density has significantly increased during plant evolution. Higher densities are observed in angiosperms compared to early land plants, and among angiosperms, recently diverged C4 species have the highest values. This has allowed leaves to increase water conductance, transpiration and possibly photosynthesis. Despite its importance, the genetic architecture of this trait is not well characterized and its relationship with photosynthesis has not been clearly established.
- Using native Mexican varieties of maize adapted to a wide range of environmental conditions, we show that vein density is variable and plastic. We leverage this variation to perform correlation analyses with photosynthetic rates and to map genetic regions associated with vein patterning traits using a MAGIC population.
- Our results show that higher vein densities are correlated with higher photosynthetic rates, but only for small intermediate veins. Varieties adapted to drier environments can substantially increase vein density in response to heat, suggesting a role in water use efficiency. We further detected 12 QTLs associated with vein patterning and identified candidate genes related to small intermediate vein development.
- These findings have implications for understanding vein architecture evolution, particularly that of C4 plants, which have significantly higher photosynthetic efficiency and productivity under warm and dry conditions.

## Introduction

Leaf venation is an important anatomical trait that significantly impacts plant performance. In particular, changes in vein density, which involve the number and spacing of veins per unit of area, are known to affect fundamental physiological features such as hydraulic conductance, gas exchange efficiency and carbon transport (Brodribb et al., 2007, Ye et al., 2021, Amiard et al., 2005), processes that influence photosynthetic rates and productivity.

Changes in vein patterning are known to have occurred during plant evolution and adaptation. For example, the emergence of early angiosperms was marked by a 3-to-4-fold increase in vein density compared to their ancestors and to other early land plants. Such an increase represented an innovation that significantly improved leaf hydraulic efficiency by decreasing the distance between the mesophyll, which moves water far less efficiently than xylem, and the site of transpiration (stomata). As a consequence, it is thought that angiosperms acquired higher transpiration rates, facilitating their early diversification (Boyce et al., 2009). This hypothesis is supported by comprehensive physiological and anatomical correlations made among representatives of extant plant clades (Brodribb et al., 2007).

Among angiosperms, a further increase in vein density occurred later in evolution during the multiple independent origin of species performing C4 photosynthesis, many of which evolved as recently as 3 mya. Indeed, increased vein density is considered to be one of the first steps towards their evolution (Lundgren et al., 2019, Sedelnikova et al., 2018, Gowik and Westhoff, 2011). In the case of C4 grasses, high vein density has been achieved through the development of smaller veins, which are classified based on their ontology: mid and lateral veins (lv) are the first to develop, followed by rank1(r1) and rank2 (r2) veins, the latter being exclusive to C4 species (Sedelnikova et al., 2018). For C4 plants, vein density is not only relevant for hydraulic conductance, but it is also hypothesized to impact photosynthetic capacity more directly by modifying the bundle sheath to mesophyll ratio in the leaf. This ratio is important because the vein-associated bundle sheath became the main site of photosynthesis in C4, as opposed to the more abundant mesophyll in their C3 ancestors.

A correlation between vein density and photosynthetic rates is thus suggested by several lines of evidence and it is certainly relevant to explain adaptation events, as well as an important breeding target. However, most of this evidence is indirect, and although the relationship between vein density and hydraulic conductance is well supported, a correlation between vein density and photosynthesis is less clear, with studies showing contrasting results (Rishmawi et al., 2017, Cohu et al., 2013, Feldman et al., 2017). This is likely due to comparisons being mostly made between individuals from different species, or mutants which also vary in fundamental properties like leaf shape, thickness, and physiology. Hence, more studies are necessary to support this hypothesis.

In addition to revealing the relationship between vein patterning and fundamental processes such as photosynthesis, defining its genetic basis is critical to understanding underlying developmental mechanisms. To date, several genetic regulators of vein patterning have been described, mainly members of the *SCARECROW (SCR)/SHORT ROOT (SHR)* pathway (Slewinski et al., 2014). Both *SCR* and *SHR* regulate cell divisions in the leaf ground tissue giving rise to the innermost mesophyll layer. Mutants show a reduced number of mesophyll cells between adjacent veins, leading to changes in vein density (Hughes et al., 2019, Liu et al., 2023). In addition, members of the *INDETERMINATE DOMAIN (IDD)* transcription factor family have recently been found to also have a role in ground tissue cell divisions by interacting with both SHR and SCR as part of a gene expression feedback loop (Hughes et al., 2023). One transcription factor named *TOO MANY LATERALS (TML)* has been found to regulate vein density independently of the *SHR/SCR* pathway. Unlike *SHR* and *SCR*, *TML* does not affect vein density by regulating mesophyll cell divisions, but by suppressing large lateral vein development (Vlad et al., 2024). Although these discoveries have allowed the field to advance forward, important questions remain to be answered. For example: are different vein types controlled by independent genetic elements? And in that case, what are the developmental regulators directly controlling the biogenesis of r2 veins, which contribute more to increased density and are especially important in C4 physiology?

In this work, we first test the hypothesis that vein density significantly contributes to photosynthetic efficiency and then use quantitative genetics to propose putative novel regulators controlling vein patterning. We selected 8 maize native Mexican varieties (NMVs) adapted to contrasting environmental conditions (mainly precipitation, temperature and altitude) and found them to show significant variation in vein density and maximum photosynthetic rates. We leveraged this diversity to correlate anatomical and physiological data, showing that increased rank 2 vein density leads to higher photosynthetic rates, especially in those species adapted to arid environments. Furthermore, we mapped genetic regions associated with vein architecture traits in a Multi-parent Advanced Generation Inter Cross (MAGIC) population, which was developed by intercrossing the 8 NMVs described above, and complemented with transcriptomic data from leaf primordia to identify a short list of candidate regulators. To our knowledge this is the first study in maize that uses linkage mapping to identify genetic regions controlling vein patterning.

## Results

### Mexican Native Maize Varieties (NMV) exhibit significant variation in vein patterning

To test the hypothesis that adaptation to a wide range of temperature and precipitation, would result in intraspecies photosynthetic phenotypic diversity, we selected 8 Native Maize Varieties (NMV) originating from contrasting environments in Mexico. These varieties are Gordo (GOR), Jala (JAL), Mushito (MUS), Nal-Tel (NAL), Reventador (REV), Palomero Toluqueño (PAL) and Zapalote Chico (ZAP). Their distribution ranges from the Peninsula de Yucatán in the south (16.3458°S) to Chihuahua in the north (29.68°N) (Fig. 1A). Importantly, along this geographic range environmental conditions vary significantly. For example, during the growing season GOR is adapted to complete its life cycle in temperatures that oscillate between 14 to 19 °C, while ZAP grows at an average of 30 °C. Precipitation can vary from a maximum of 160 mm for GOR to more than 400 mm for REV and altitudes range from 0 (NAL) to 2600 meters above sea level (PAL) (Fig. 1A & C). We obtained the aridity index (AI), evapotranspiration (PET), and seasonal accumulated precipitation (AP) for the source environment of each variety and performed principal component analyses (PCA). Environmental PCA shows that GOR and ZAP source environments are the most positively correlated with aridity index (AI) and potential evapotranspiration (PET), while NAL and REV source environments are the most correlated with mean temperature (MT) and seasonal accumulated precipitation (AP) (Fig. 1B).

**Figure 1.**
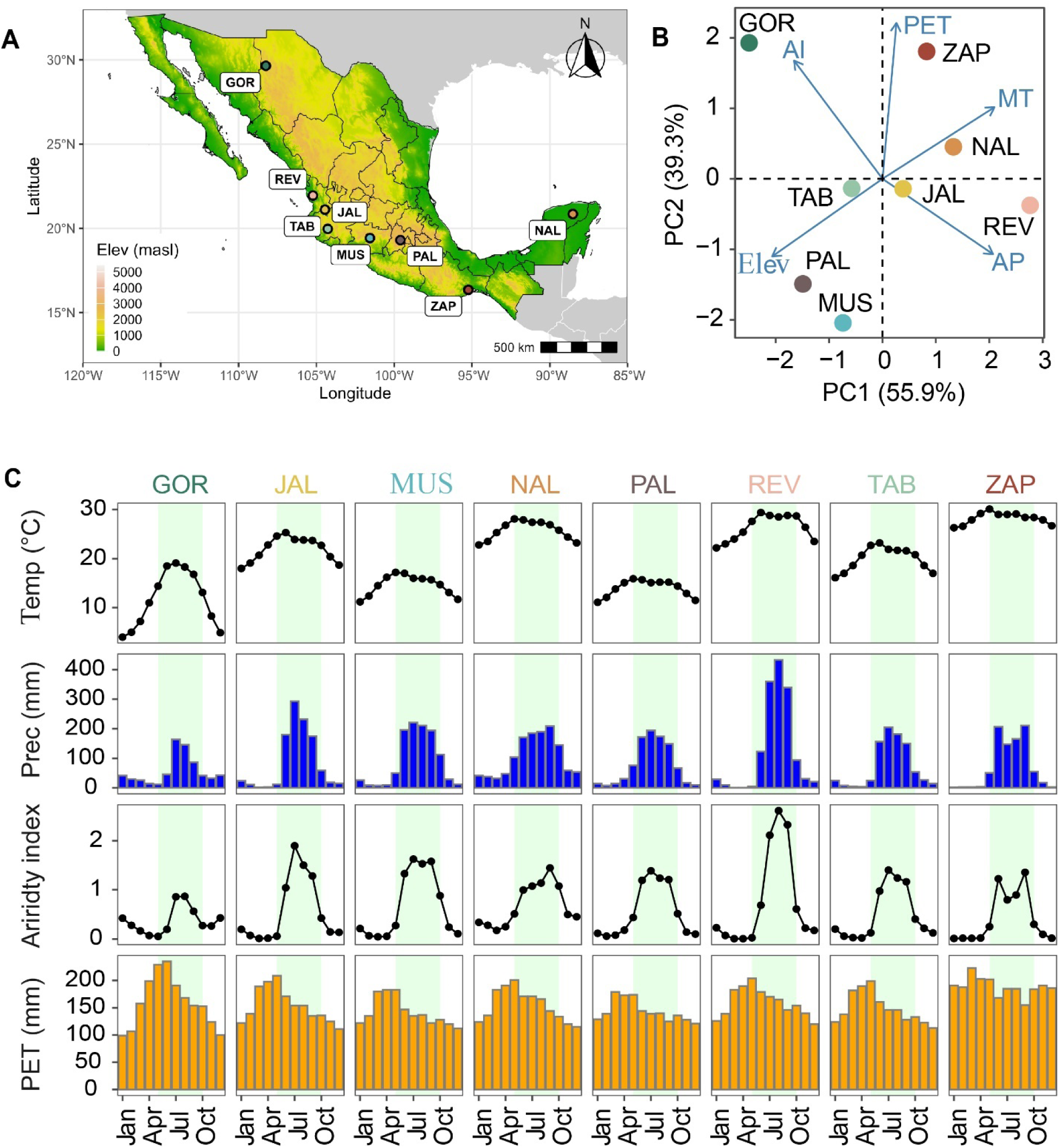
Environmental variables associated with NMVs geographic origin. (**A)** Geographic origin of each NMV. The map is colored according to elevation scale. (**B)** Principal component analysis (PCA) of environmental variables associated with NMVs geographic origins, including elevation (Elev), seasonal mean temperature (MT), seasonal potential evapotranspiration (PET), Seasonal Aridity Index (AI), and seasonal accumulated precipitation (AP). The biplot shows PC1 explaining 55.9% variance and PC2 explaining 39.3% variance. (**C)** Dynamics of main environmental variables associated with each NMV throughout the year. The green bar indicates the crop’s growing season (May to Oct).

To assess differences in vein patterning in the NMVs, we developed a method to prepare histological sections based on manual transverse dissections of leaves embedded in agar, followed by staining with a fluorescent dye (Fig. S1). Samples were collected and prepared at Vegetative Stage 5 (V5) and r1, r2, and lateral vein densities quantified (Fig. 2A). We found phenotypic variation in all anatomical traits measured. r2 vein density (r2vd) ranged from 39.05±1.5 veins/cm in GOR to 84.03±0.30 in JAL (Fig. 2B). r1 vein density (r1vd) varied from 4.05 ± 0.20 to 12.0 ± 0.30 veins/cm (Fig. 2C), while lateral vein density (lvd) varied from 3.5 ± 0.25 to 4.5 ± 0.15 veins/cm (Fig. 2E). We further calculated the ratio of r1 to r2 veins (r1/r2) to determine if there were differences in the proportion of small intermediate veins across varieties. Low values indicate a higher number of r2 veins in relation to r1 veins. Values ranged from 0.28 ± 0.02 (GOR) to 0.05 ± 0.003 (JAL), a five-fold difference (Fig. 2D). Therefore, varieties from the same species and photosynthetic type adapted to contrasting environmental conditions can show considerable differences in vein morphological traits.

**Figure 2.**
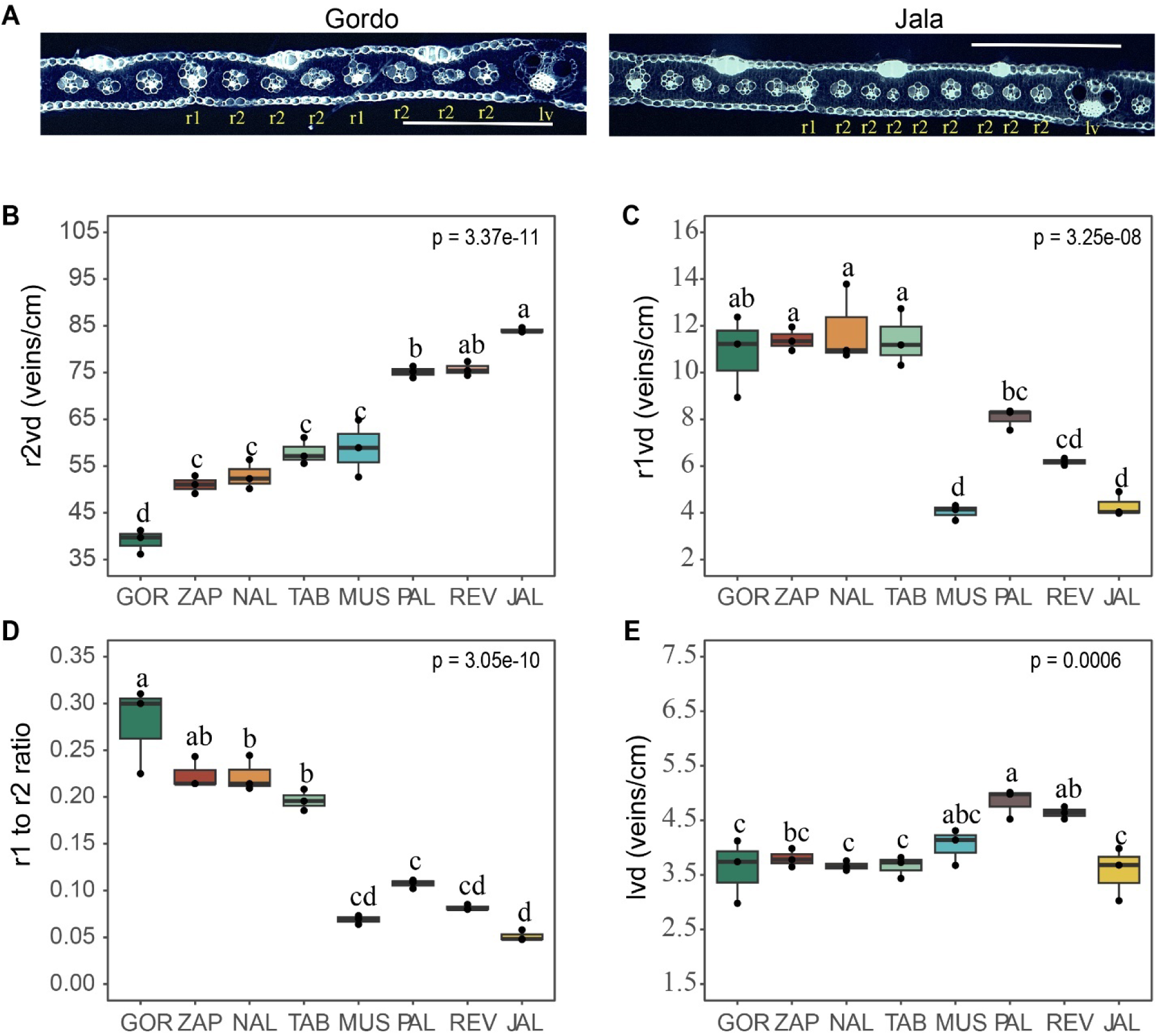
Phenotypic variation of leaf venation traits across NMVs. (**A)** Representative images of leaf cross sections from Gordo and Jala varieties, representing extreme variation phenotypes. Vein type classification is shown as Rank1 (r1), Rank2 (r2) and lateral veins (lv) below each vein. The upper side of the leaf in the image is the adaxial surface. (**B)** Quantification of r2 veins/cm across varieties. **(C)** Quantification of r1 veins/cm across varieties. **(D)** Quantification of r1 to r2 ratio across varieties. **(E)** Quantification of lv veins/cm across varieties. ANOVA was applied to determine significant differences between varieties. Letters above individual box-scatterplots indicate significant groupings according to Tukey’s means comparison (*α* = 0.05). Three replicates per variety were included. Bars = 500 mm (A). Bold bar in boxplots represents the median.

### NMVs exhibit ontogenic plasticity for leaf vein density as a function of temperature

Our initial assessment of vein density was performed on plants grown in a warm greenhouse with an average temperature of 30 °C. However, some native varieties are naturally adapted to cooler climates and growing at higher temperatures may induce developmental changes and affect vein density estimates. Indeed, some studies have demonstrated plasticity of vein traits in natural populations of a given species exposed to different environmental conditions (Garthwaite et al., 2024, von Arx et al., 2012).

Therefore, we evaluated developmental plasticity for vein density in response to changes in temperature. We grew our NMV in parallel under two conditions: a warm and a cool greenhouse, at an average temperature of 30 °C and 20 °C respectively. We collected tissue at two developmental time points: V5, leaf 5 and V9, leaf 9. Since it is known that maize seeds already contain four to five fully formed embryonic leaves (Liu et al., 2013), we hypothesized that vein patterning in the fifth leaf would already be genetically determined and thus not reflect environmental changes, representing a developmental base line. In contrast, the 9^th^ leaf collected at V9 should reflect any developmental adaptations in response to temperature.

We observed that most varieties significantly increased vein density from V5 to V9 when grown in a warm environment, but not when grown under cooler conditions (Fig. 3A). Notably, this response was observed for r2 veins, but not for r1 or lateral veins (Fig. S2). When comparing vein density from leaf 5 (V5) within varieties there was no significant change across environments, indeed suggesting that development of the fifth leaf was already determined from the seed stage. In contrast, for most varieties r2 vein density measured at leaf 9 changed across treatment (Fig. 3A). For example, vein density for GOR at leaf 9 increased from approximately 55 veins/cm at 20 °C to almost 70 veins/cm when grown at 30 °C. Similarly, NAL, MUS, and JAL showed a significant increase. Notably, not all varieties show the same response. Vein density did not change in ZAP at V9 across conditions, and TAB even showed a reduction in vein density under the warm environment (Fig. 3A, upper panel). We plotted phenotype variation across varieties as a function of temperature to visualize plasticity norms of reaction and found several types: genotype but no environment effect (Fig.3B, left), both genotype and environment effect (Fig.3B, center), and genotype per environment interaction effect (Fig.3B, right), showing that evaluated varieties expressed different phenotypic ranges in response to temperature.

**Figure 3.**
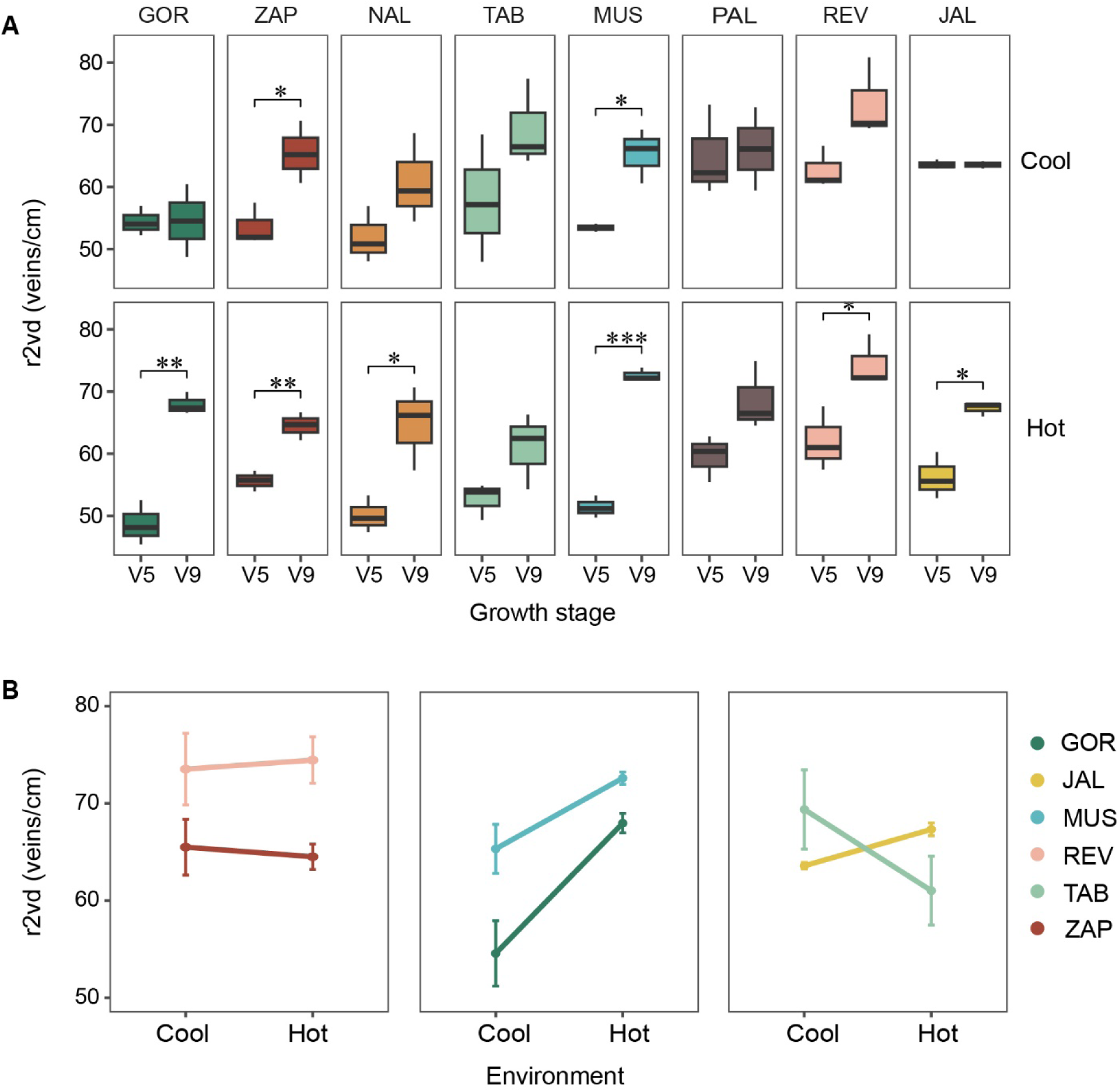
Changes in vein density across development and temperature. (**A)** Quantification of r2 vein density at developmental stages V5, leaf 5 and V9, leaf 9 in two environmental conditions (Cool, 20°C; warm, 30°C). (**B)** r2 vein density changes in response to temperature measured at V9, leaf 9 showing reaction norms. Genotypes were plotted in different panels for clarity.

### Small intermediate vein density is correlated with CO_2_ assimilation rates

Having found significant variation in vein architecture traits across our NMVs, we set out to investigate how these differences would correlate with photosynthetic capacity. We observed that CO_2_ assimilation curves (A/Ci) differed significantly across varieties and environments (Fig.4A, left). Differences in maximum CO2 assimilation (Amax) ranged from 12 µmol m^-2^ s^-1^ in NAL to 22 µmol m^-2^ s^-1^ in GOR under cold conditions, and from 28 to 45 m^-2^ s^-1^ in JAL and ZAP respectively under warm conditions. Similarly to the responses observed for vein density, most varieties showed a significant increase in Amax under warm conditions when measuring photosynthesis in leaf 9 (Fig.4A). Moreover, we noticed that some varieties had a similar response both in vein density and photosynthesis. For example, NAL shows an increment in Amax and r2vd in leaf 9 only at 30 °C, but not when grown at 20 °C (Fig. 4A, right).

**Figure 4.**
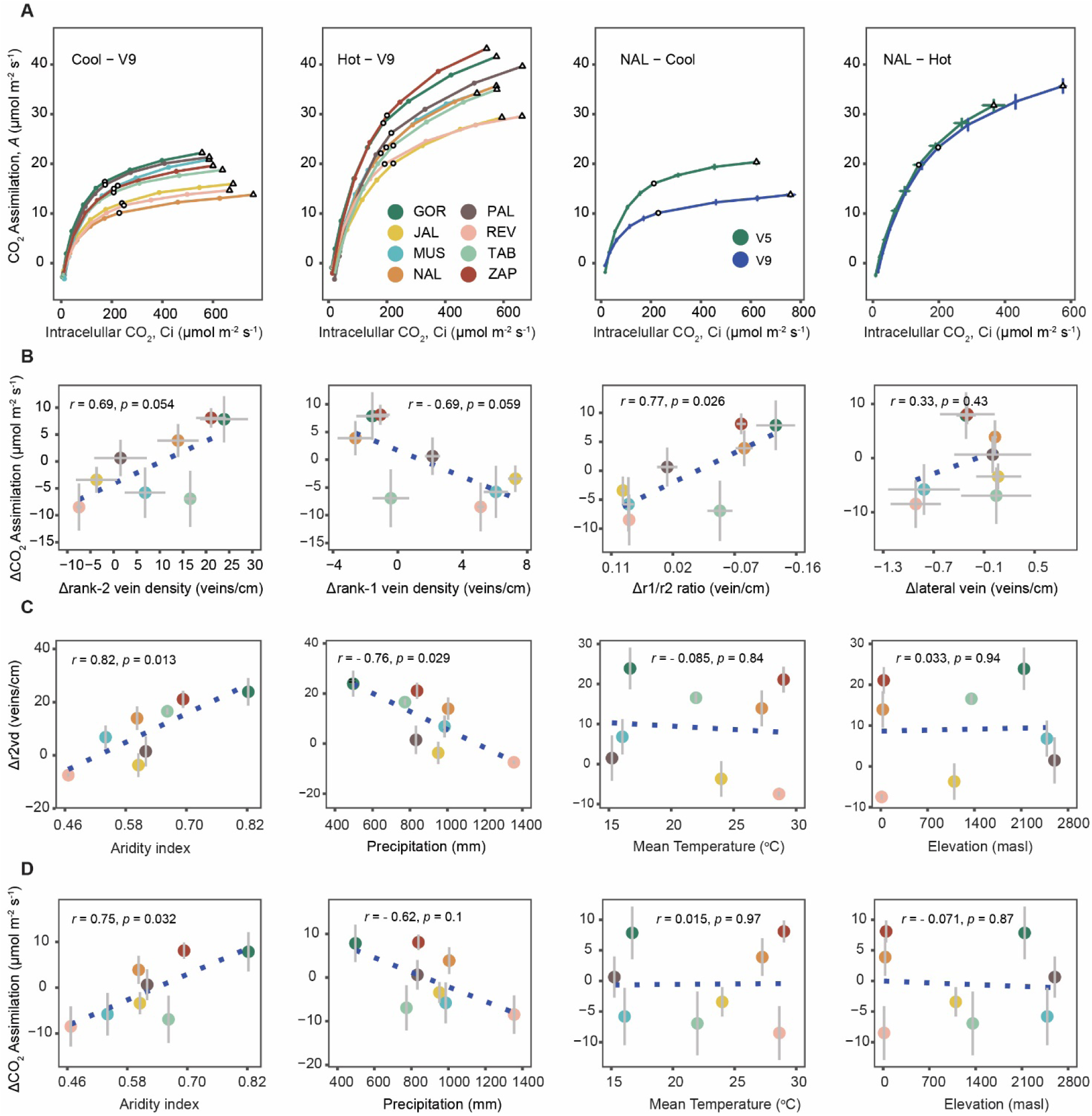
Correlation between CO_2_ assimilation, anatomical and environmental variables. (**A)** A-Ci curves measured in two environmental conditions (Cool and Warm) for NMVs at V9 Stage (leaf 9), and for NAL across developmental stages in Cool and Hot conditions. (**B**) Scatterplots showing the relationship between changes in CO_2_ assimilation and changes in vein density traits across NMVs. (**C**) Scatterplots showing the relationship between changes in vein density and environmental variables collected at the NMVs geographic origins. (**D**) Scatterplots showing the relationship between changes in CO_2_ assimilation and environmental variables collected at the NMVs geographic origins. Pearson correlation coefficients are shown at the top.

Photosynthetic rates depend on several anatomical (other than vein density) and physiological factors, including leaf area and thickness, chlorophyll content, and stomatal density (Wang et al., 2016, Harrison et al., 2020, Hu et al., 2020). Hence, variation in these traits across NMVs tested could contribute to differences in Amax that would be unrelated to vein density changes. To assess the possible effect of such confounding variables, we first quantified variation in those traits across genotypes. We found variation for leaf area, stomatal density and relative chlorophyll content. Only leaf thickness was constant (Fig. S3A-D). Next, we assessed if these traits were also plastic (i.e. if they changed from V5 to V9) and found that, unlike vein density, there was no change across development, except for leaf area (Fig. S3E-H). However, the magnitude of change for the latter trait was similar across all genotypes, probably reflecting a normal increase in leaf size as plants grew older.

Hence, we leveraged the fact that NMVs exhibit different degrees of phenotypic change (plasticity) for vein density and Amax, not observed in other measured traits, to assess correlation between the magnitude of change for both variables. Thus, minimizing the effect of confounders due to inter-genotypic morphological variation. We calculated the change in vein density (Δvd) and the change in maximum CO_2_ assimilation rate (ΔAmax) from V5 to V9 and include those variables in our correlation analyses. We first explored the best predictor variables for a model in which Amax or ΔAmax were the single dependent variables. We used a bidirectional linear stepwise regression model including 12 vein density traits as variables: absolute values from V5 and V9, as well as Δvd, for all vein types across varieties (Table S1). No significant predictors were found when using absolute Amax values; however, a best model was found for ΔAmax in which the change in r1 to r2 vein ratio (Δr1/r2) was the best significant predictor (r^2^ = 0.60, p = 0,025). This suggested that changes in minor vein density from V5 to V9 were strongly associated with maximum photosynthetic rates.

We further calculated the Pearson correlation coefficient for individual comparisons and found that Δr2vd is positively correlated with ΔAmax (r = 0.69, p = 0.05). In contrast, Δr1vd is negatively correlated with ΔAmax (r = 0.69, p = 0.059) and Δlvd does not show a significant correlation (r = 0.33, p = 0.43). As expected from the model prediction, the strongest correlation was found between Δr1/2 vein ratio and ΔAmax (r = 0.77, p = 0.026) (Fig. 4B). Hence, these results indicate that vein density, specifically r2 veins, strongly influences photosynthetic rates, and that other vein types like r1 and lv seem to be less important for maintaining photosynthetic capacity. Additionally, we assessed correlation between ΔAmax and Δleaf area, since this variable also changed from V5 to V9, and found no significant correlation (*r* = -0.18, *p* = 0.67), suggesting leaf area does not influence the association between our two phenotypes of interest (Fig. S4A).

It has been suggested that increased vein density is an adaptation to dry and warm environments, although previous studies show that plants growing in these conditions do not always have higher vein densities, at least when comparing absolute values (Garthwaite et al., 2024, Wang et al., 2024). Following this hypothesis, we examined the correlation between Δr2vd and environmental variables collected from the source environment of each variety. We observed a correlation between Δr2vd and both precipitation and aridity (Fig. 4C), such that varieties sourced from drier environments more strongly increase vein density through development when exposed to heat, even though they may not have the highest absolute values initially. In contrast, varieties sourced from humid conditions did not significantly increase vein density under warm conditions, and in some cases vein density even decreased at V9. No significant correlation was found between r2vd absolute values and aridity or temperature.

We did not find a significant correlation between Δr2vd and temperature or elevation (Fig. 4C, right). Therefore, vein density plasticity is correlated with heat but mainly in varieties sourced from arid environments. Similarly, we found that maximum photosynthetic capacity is also correlated with precipitation and aridity but not to temperature and elevation (Fig. 4D).

### The MexMAGIC population reveals genetic regions associated with vein traits

We showed that vein patterning is important for maximum photosynthetic capacity and that increased r2 vein density may be relevant for adaptation to arid environments. However, the genetic basis of this trait is still not well defined, and it is not clear if different genetic pathways control development of individual vein types. Hence, we employed an eight-founder multi-parent advanced generation intercross (MAGIC) population developed from the same NMV described above to map genetic regions associated with vein patterning. The eight founders were intercrossed for three generations followed by three rounds of plant intermating and three rounds of self-pollination (“I3S3”) (Fig. 5A). Over 240 MAGIC families were produced, genotyped and evaluated in the field under similar conditions as reported for our warm greenhouse experiments.

**Figure 5.**
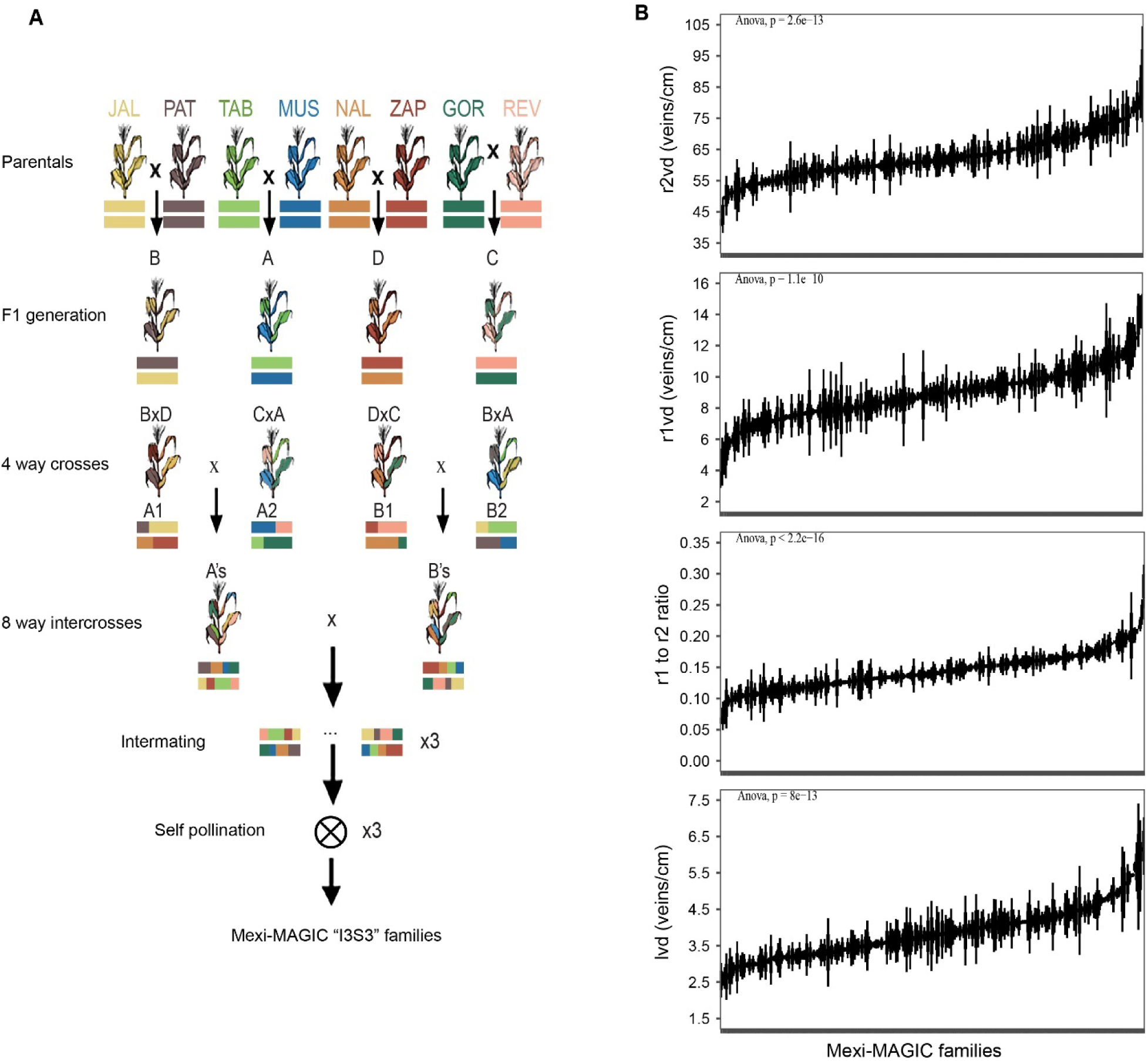
Quantification of vein density traits across the MexMAGIC population. **(A)** Crossing scheme for the generation of the MexMAGIC population from the 8 NMVs. (**B**) Quantification of r1, r2, r1 to r2 ratio, and lateral vein density across 240 families from the MexMAGIC population.

We employed our rapid histology technique to image leaf cross sections and evaluate r2, r1, and lateral vein density, as well as r1 to r2 vein ratio in 480 individuals (duplicates per family). We found that trait variation in the field-grown MAGIC population was continuous, suggesting it is indeed quantitative, and that it recapitulated that of the parentals (Fig. 5B). In addition, broad sense trait heritability was high, ranging from h^2^ = 0.58 for r1vd to 0.73 for r1 to r2 ratio, showing a substantial contribution of genetic factors to trait variation (Table S2).

### Twelve QTLs are related to vein architecture traits in the MexMAGIC population

We integrated both QTL and GWAS analysis to map genetic regions associated with vein traits. A set of ∼1M SNPs distributed across the 10 maize chromosomes were used as markers for the analysis (Table 1). We identified 12 QTLs for vein density traits. Only those loci that showed a significant association in both models, QTLs (alpha = 0.05, 1000 permutations) and GWAS were retained (Fig. 6). QTL regions were defined as the regions above a LOD score of 4.87. Notably, different QTLs were associated with each vein type trait (e.g. there were no overlapping regions for r1, r2 and lv), thus suggesting they have different genetic architectures. This is also supported by the low correlation observed between phenotypic traits (Fig. S4B). Of particular interest were genetic regions controlling development of r2 veins, since we showed that this vein type is especially important for photosynthetic capacity. We found one QTL in chromosome 1 (qr2vd_1_1) that explained 13% of r2vd variance, and three QTLs in chromosome 8 (qr2vd_8_1, qr2vd_8_2, and qr2vd_8_3) that together explained 40.2% of phenotypic variance for r2vd (Fig. 6). Although our QTL call criteria identified three regions in chromosome 8, it is also possible that these represent a single locus. The physical size of the mapped regions ranged from 6.9 megabases (Mb) in the case of the qr2vd_8_3 QTL to 17.8 Mb for qr2vd_8_1 (Table 2).

**Figure 6.**
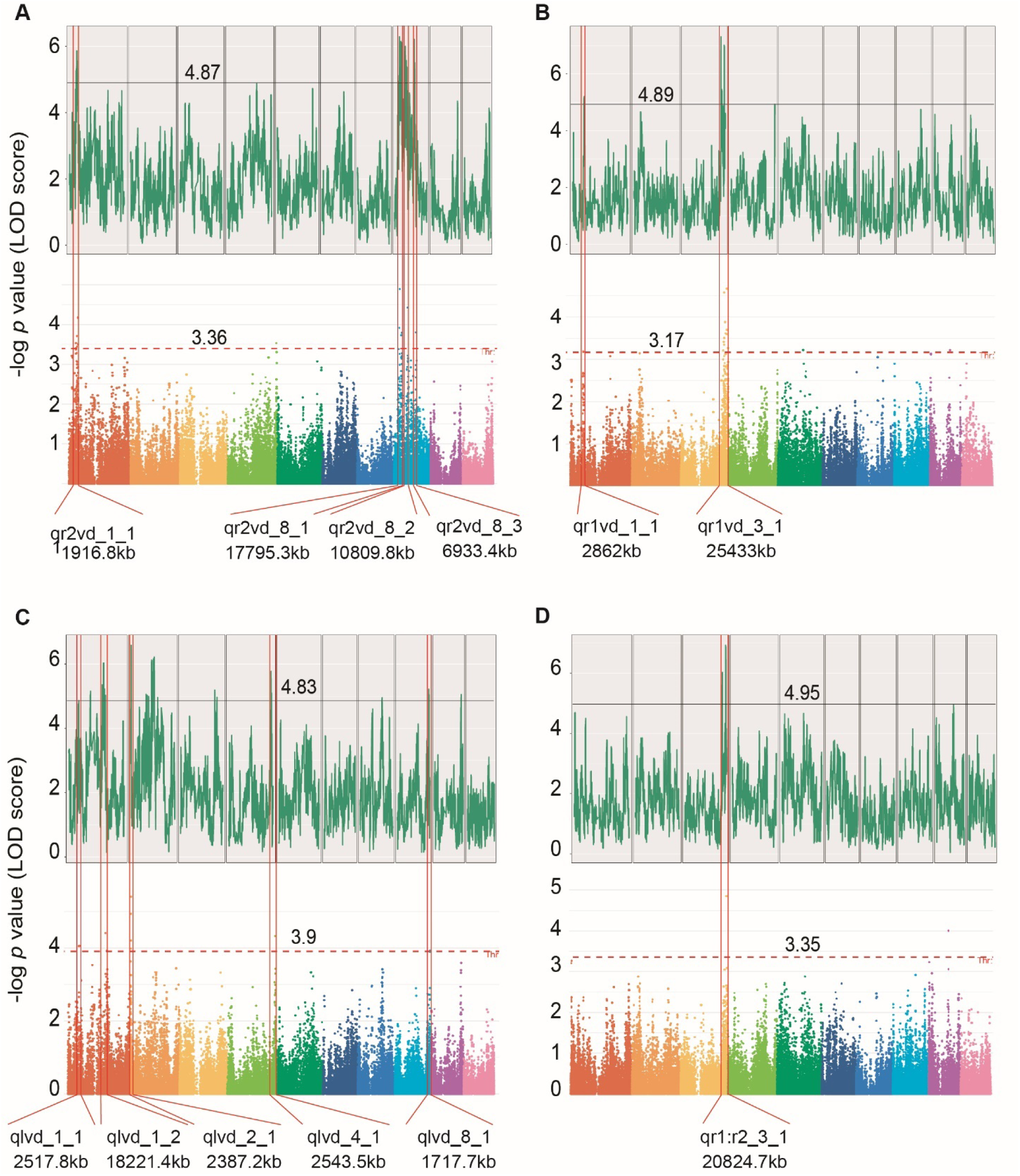
Overlapping genetic regions identified through QTL and GWAS association models. **(A-D)** Test statistic plot -Log10(p) against genome location (up) and Manhattan plot showing association for r2, r1, lateral vein density and r1 to r2 ratio. The dotted lines show the critical value of the test statistic from 1,000 permutations (p < 0.05). Values showing the size of the overlapping regions are indicated below each panel. Colors indicate different Chromosomes. Lines across panels indicate overlapping regions from QTL and GWAS models.

**Table 1.** Genome-wide distribution of 1M SNPs markers across 10 chromosomes used for linkage mapping in the MexMAGIC population.

| Chr | SNPs | % SNP | Lo lim (bp) | Up lim (bp) | Dist (Mb) | Dens (SNPs/Mb) |
| --- | --- | --- | --- | --- | --- | --- |
| 1 | 164,807 | 16.48 | 26,435 | 306,781,408 | 306.75 | 537.26 |
| 2 | 91,406 | 9.14 | 40,232 | 244,371,401 | 244.33 | 374.11 |
| 3 | 121,087 | 12.11 | 91,830 | 235,604,954 | 235.51 | 514.14 |
| 4 | 130,728 | 13.07 | 135,286 | 246,943,111 | 246.81 | 529.68 |
| 5 | 106,358 | 10.64 | 11,836 | 223,714,157 | 223.70 | 475.44 |
| 6 | 83,458 | 8.35 | 114,804 | 173,322,741 | 173.21 | 481.84 |
| 7 | 68,894 | 6.89 | 95,329 | 182,153,951 | 182.06 | 378.42 |
| 8 | 93,715 | 9.37 | 71,443 | 181,085,133 | 181.01 | 517.72 |
| 9 | 73,548 | 7.36 | 16,172 | 159,687,184 | 159.67 | 460.62 |
| 10 | 65,919 | 6.59 | 127,325 | 150,830,223 | 150.70 | 437.41 |
| Sum | 999,920 | 100 |  |  | 2103.76 | 4706.64 |

**Table 2.**
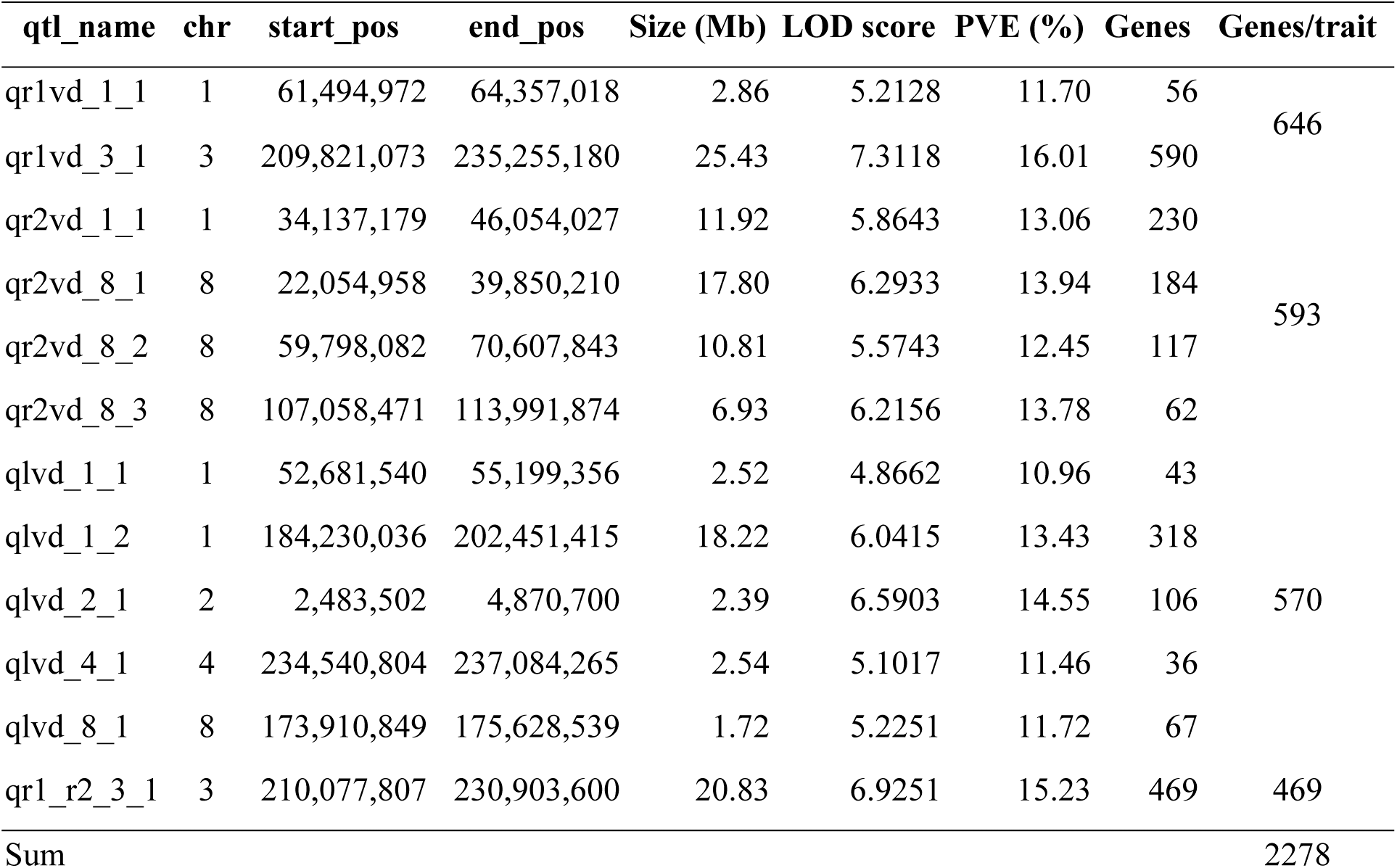
Information on the size, position, and coding genes contained in the identified QTLs for vein density traits.

Regarding r1 veins, we detected two QTLs, one in chromosome 1 (qr1vd_1_1) that explained 11.7% of r1vd phenotypic variance, and another in chromosome 3 (qr1vd_3_1) that explained 16% of the variance. The latter was the largest detected region overlapping with a single QTL for r1 to r2 ratio, suggesting that association between these traits was mostly influenced by variation in r1 veins (Fig. 6, lower right). Finally, five QTLs were associated with lateral vein density. Two in chromosome 1, and single QTLs in chromosome 2, 4 and 8, which explained 11%, 13.5%, 14.5%, 11.5%, and 11.7 of the lvd phenotypic variance respectively (Table 2). QTLs coordinates and genes contained in each region are detailed in Table S3. Next, we determined gene expression profiles in leaf primordia to narrow down a list of probable candidate genes regulating these traits.

### Identification of putative regulators of r2 vein development through transcriptome to phenotype correlation analyses

To narrow down the number of candidate genes for vein traits, we generated transcriptomes from early leaf development for all maize varieties. It is known from previous studies that veins begin to form early during leaf development. In particular, r2 veins are specified during plastochron 4 (Perico et al., 2024, Robil and McSteen, 2023). Therefore, genes controlling this trait are expected to be expressed at relatively high levels during this stage. We grew the eight founder varieties under warm conditions and dissected leaf primordia at plastochron 4 to generate transcriptomes both at V5 and V9 stages. We first assessed the expression of genes overlapping with QTL regions and retained only those showing a significant expression (Fig. 7A). After applying this filter, approximately 50% of genes were retained for further analyses (Table S4). We next assessed the expression pattern of selected genes across varieties and developmental stages (Fig.7b) and identified those whose change in expression from V5 to V9 (Δexp) showed either positive or negative correlation with Δvein density (from V5 to V9) across varieties (Fig. 7B & C). We report the top 20 genes based on Pearson correlation coefficients (*r* > 0.7, *p* = ≤ 0.05) (Fig.7D, Table S4). To further support candidate selection, we then calculated the parental genetic effects (BLUP effects for r2vd) for leading alleles (LOD ⋝ 4) located at each of the 20 genes and followed two strategies: first assess correlation between allele effects and a set of environmental variables related with temperature and humidity (Fig. 7E); and second, calculate correlation between parental effects and gene expression levels (Fig. 7F).

**Figure 7.**
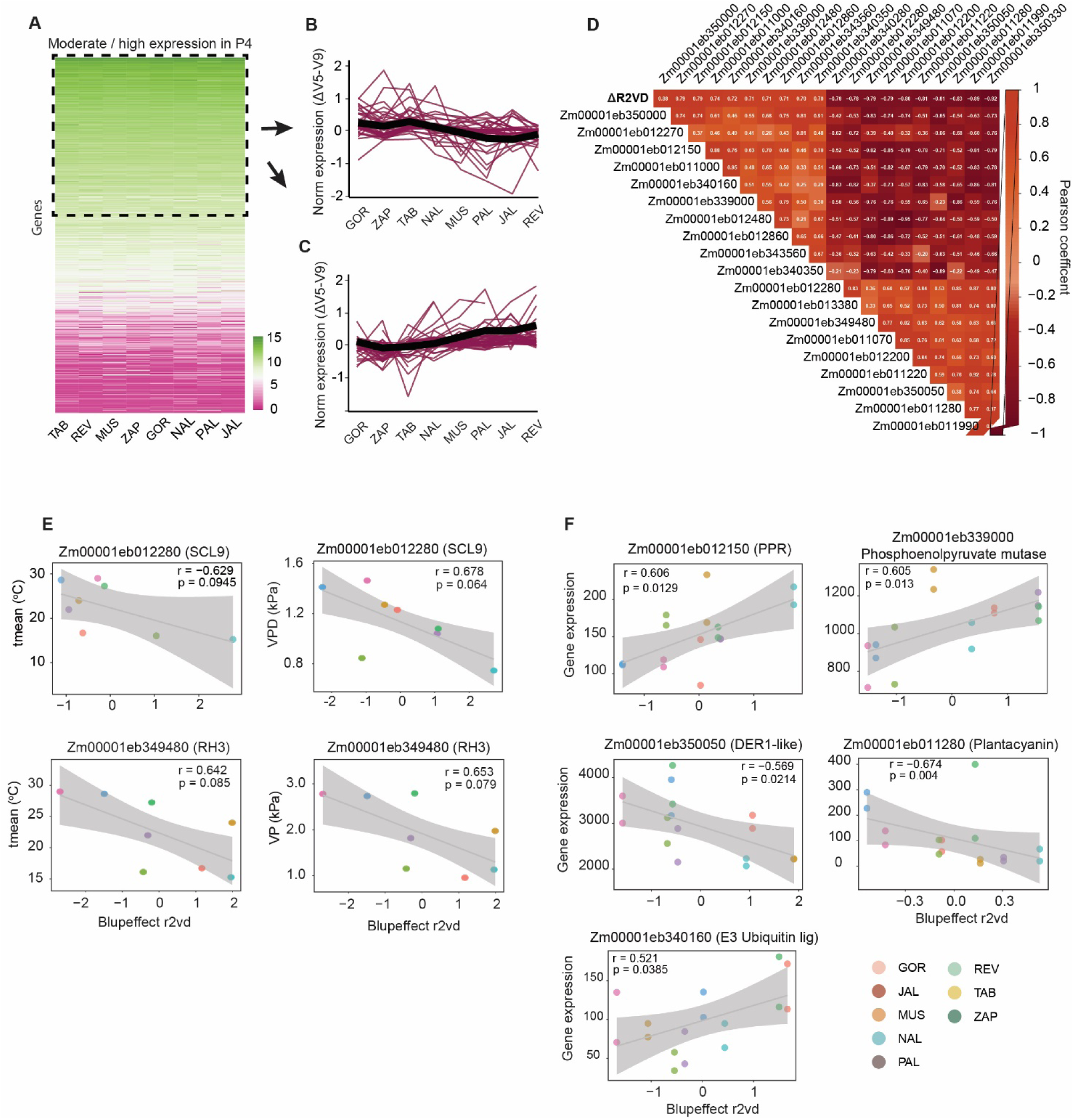
Correlation between changes in gene expression and vein density across NMVs. **A)** Heatmap representing the normalized expression values from plastochron 4 of the 568 coding genes contained in QTLs for r2 vein density. Genes with moderate to high expression are highlighted. (**B**) Expression profiles of genes (ΔV5-V9) showing a positive correlation with vein density changes (r > 0.5) across NMVs. **(C)** Expression profiles of genes (ΔV5-V9) showing a negative correlation with vein density changes (r < -0.5) across NMVs. The black line indicates the expression trend. **(D)** Correlogram showing the top 10 genes IDs with the highest positive and negative Pearson correlation coefficient between changes in expression and changes in vein density (p < 0.05). **(E)** Correlation between parental genetic effects (BLUP effects for r2vd) for leading alleles located at genes shown in (D) and source environmental variables. Correlations > *r* = 0.5 are shown. **(F)** Correlation between parental genetic effects (BLUP effects for r2vd) for leading alleles located at genes shown in (D) and gene expression levels. Correlations > *r* = 0.5 are shown.

We found allele effects for two genes that were correlated with source environmental variables. A near clinal effect was found between r2vd and both vapor pressure deficit (vpd) and temperature for the *SCARECROW-LIKE* transcription factor *Scl9* (*Zm00001eb012280*). A similar association was found between the allele effects of the DEAD-box RNA helicase *RH3*, (*Zm00001eb349480*) and source environmental vapor pressure (evp), and temperature (Fig. 7E). Both vpd and evp are determined by the amount of humidity in the environment and predict plant transpiration and desiccation. Notably, members of the *SHORTROOT/SCARECROW* family of transcription factors have been implicated in vein patterning in maize and *Arabidopsis* (Hughes et al., 2023, Hughes et al., 2019). On the other hand, homologues of RH3 are involved in intron splicing and small RNA loading in chloroplasts.

Additionally, we identified allele effects on five genes showing correlation with gene expression levels from leaf primordia (P4) (Fig. 7E): A Plantacyanin-like gene (*Zm00001eb011280*) from the qr2vd_1_1 region, with homologs functioning in light mediated regulation of hormone balance and development in *Arabidopsis* (Yang et al., 2024, Jiang et al., 2021); a pentatricopeptide repeat gene (PPR, *Zm00001eb012150*) also from the qr2vd_1_1 region, which is member of a large family of RNA-binding proteins involved in chloroplast RNA splicing; a RING-type E3 ubiquitin ligase (*Zm00001eb340160*) from the qr2vd_8_1 region; a phosphoenolpyruvate mutase (*Zm00001eb339000*) from qr2vd_8_1 region, and a DER1-like gene (*Zm00001eb350050*) part of the ERAD protein degradation pathway from the qr2vd_8_3 region.

Genes overlapping with the qr2vd_8_2 QTL did not show a correlation between allele effects and either environmental or expression values. Overall, using this strategy we managed to significantly narrow down a list of putative regulators of r2 vein development for future functional studies.

## Discussion

Understanding the genetic basis of vein development and the effect of venation architecture on photosynthetic efficiency is important for improving plant productivity. Historically, this task has been hindered by the lack of biological models showing variability in vein architecture and photosynthesis traits needed to allow correlation analyses and forward genetics approaches. We report that intraspecies variation exists in maize, a staple crop globally and a well-established experimental model. By measuring both vein density and CO_2_ assimilation rates in eight native Mexican varieties across temperate and warm conditions we discovered these traits are plastic. Vein density increases during development in most varieties in response to heat. This is in line with a previous study in *A. thaliana* in which plants respond to short term heat by increasing vein density and vein connections (Dhakal et al., 2021). However, other studies in dicotyledonous species have shown contrasting results and simultaneous photosynthesis measurements have not been taken to establish a correlation between both traits (Garthwaite et al., 2024). We found that such correlation indeed exists in maize, but only for r2 veins. We also found that varieties showing more plasticity are those sourced from environments with lower precipitation. This could explain the contrasting results obtained by previous studies if genotypes reflecting different adaptation histories were included.

Our findings are also relevant in the context of understanding Kranz anatomy evolution, characterized by a significant increase in vein density. Although it is generally believed that heat and a decrease in atmospheric CO2 concentrations were the major factors influencing this process, we did not observe a correlation between increased vein density and temperature of origin, nor between increased CO2 assimilation and temperature of origin. Instead, our data shows a strong positive correlation of both variables with precipitation and aridity. This supports an alternative model that considers the combined influence of multiple factors (e.g., high light, low CO_2_, heat, and aridity) to explain the emergence of C4 photosynthesis and Kranz anatomy, not only as a strategy to increase photosynthetic efficiency, but also as a water conservation mechanism that allowed these plants to dominate arid environments (Osborne and Sack, 2012).

Under the current context of climate change, there is a growing interest in characterizing the genetic architecture of Kranz anatomy to allow engineering these traits in C3 species (Karki et al., 2013). The intraspecific variation in CO_2_ assimilation and leaf anatomical traits found across native Mexican maize makes it an ideal model for identifying the genetic basis of C4 photosynthesis using quantitative genetics. Recently, a couple of studies have shown the feasibility of applying QTL mapping for C4 traits. The first takes advantage of the large variability found in the C4 dicot *Gynandropsis gynandra* to perform linkage mapping for agronomic and leaf anatomical traits (Simpson et al., 2025). In this study, one QTL for vein density was confidently mapped. The second is a genome-wide association study (GWAS) using individuals from the grass *Alloteropsis semialata*, which is the only described monocot with C3, C3+C4 intermediate, and C4 populations. Although vein density was not reported, the authors identified two genomic regions associated with bundle sheath distance, a related trait, and identified one annotated gene: *Glucan Synthase-like 8 (GSL8)*, which is known for its role in leaf development in *Arabidopsis* (Alenazi et al., 2024).

In this study, we identified 12 QTLs for vein density traits using a maize MAGIC population. Notably, we identified and quantified vein types individually, that is r1, r2 and lv, providing a high resolution in mapped vein traits. We showed that each vein type is controlled by different genetic architectures. This is relevant in the context of engineering C3 crops with increased vein density, because it suggests that r2 vein specific regulators can be targeted to improve productivity without affecting development of other conserved vein types. In this regard, we generated a short list of putative r2 vein developmental regulators by applying transcriptome to phenotype correlation analyses. We further supported candidate gene selection by assessing correlation between parental allele effects at each gene leading SNP and environmental and gene expression values. Seven genes with a large association were highlighted:

A *SCARECROW-like* gene (*SCL9*), belonging to the GRAS family of transcription factors. Members of this family, mainly *SCARECROW* and *SHORTROOT*, have been implicated in regulating patterning in roots and shoots and *SCR* has a role in Kranz anatomy development in maize (Hughes et al., 2023, Hughes et al., 2019). Although that locus was not identified in our analyses, the finding of a *SCR-like* gene suggests that different members of this family may be implicated in divergent roles in vein development and patterning. Importantly, a *SCL9* gene was associated with the strength of the C4 cycle (δ13C) in an independent GWAS study conducted using *Alloteropsis semialata* populations (Alenazi et al., 2024), indicating these homologues may be important for C4 photosynthesis across grasses.

From our short list of candidates, several genes located in different QTLs are involved in post-transcriptional and post-translational modification. For example, two genes which have homologs in other species with a role in gene splicing (PPR and RH3); as well as two genes involved in protein degradation and homeostasis (DER1 and an E3 ubiquitin ligase). Hence, post-transcriptional (mainly gene splicing) and post-translational modification may be overlooked mechanisms modulating the differential action of developmental regulators from C3 and C4 species.

## Conclusion

We show that vein development is plastic in some maize genotypes, with varieties adapted to arid environments being able to substantially increase vein density in response to high temperatures. We also provide evidence that high vein density increases photosynthetic capacity and identified 12 QTLs associated with this trait. We combined transcriptomic analyses to present a shortlist of candidate genes regulating r2 vein development. These included genes with homologs involved in developmental patterning, gene splicing and post-translational regulation, which have not been characterized in this context before. Overall, our findings have implications for understanding the evolution of photosynthetic efficiency and for engineering C4 traits in C3 crops like rice and wheat.

## Supplemental Figures

**Fig. S1.**
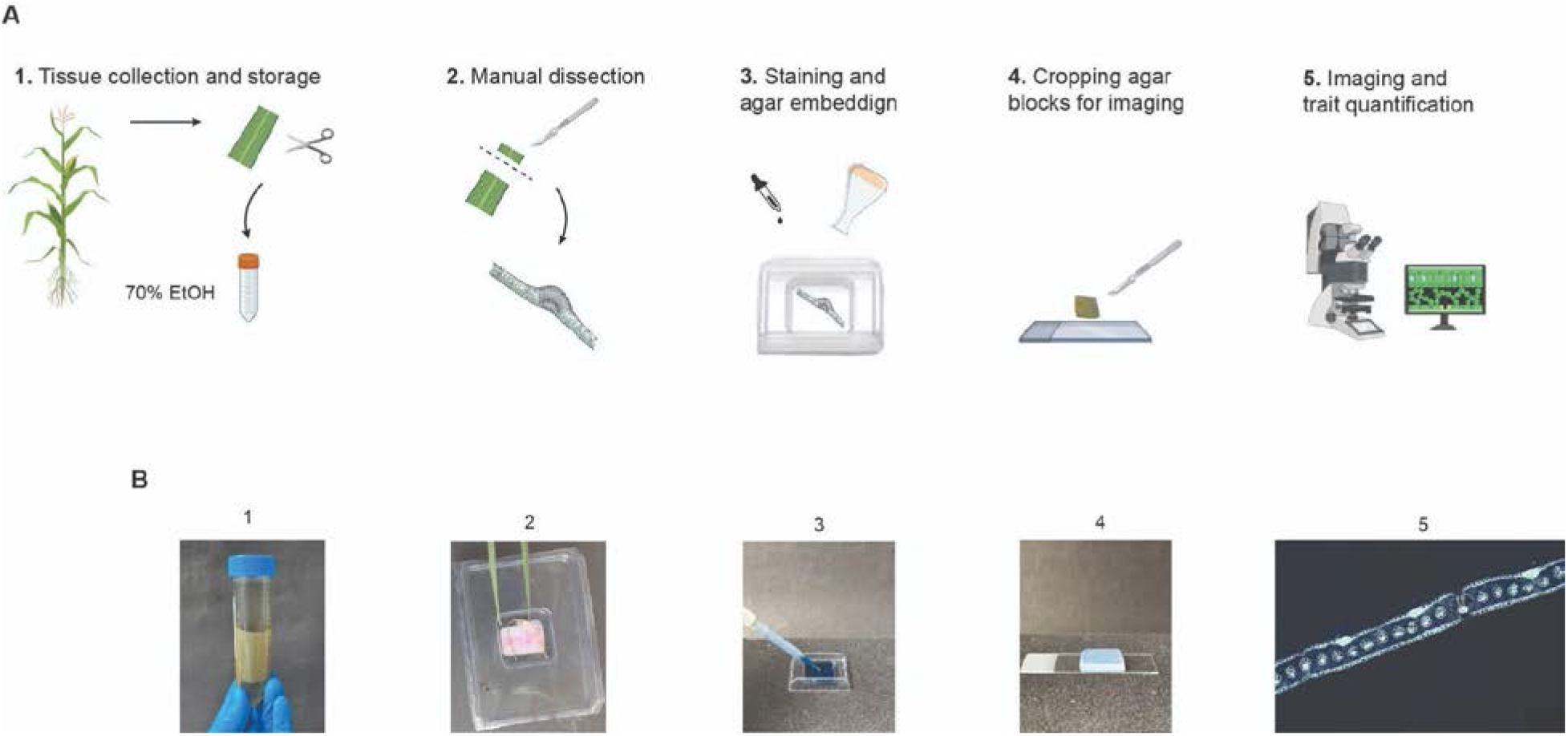
Rapid leaf histology protocol workflow. **(A)** Diagram illustrating the main steps of the rapid leaf histology protocol. **(B)** Representative images of the same main steps presented in (A) for reference purposes.

**Fig. S2.**
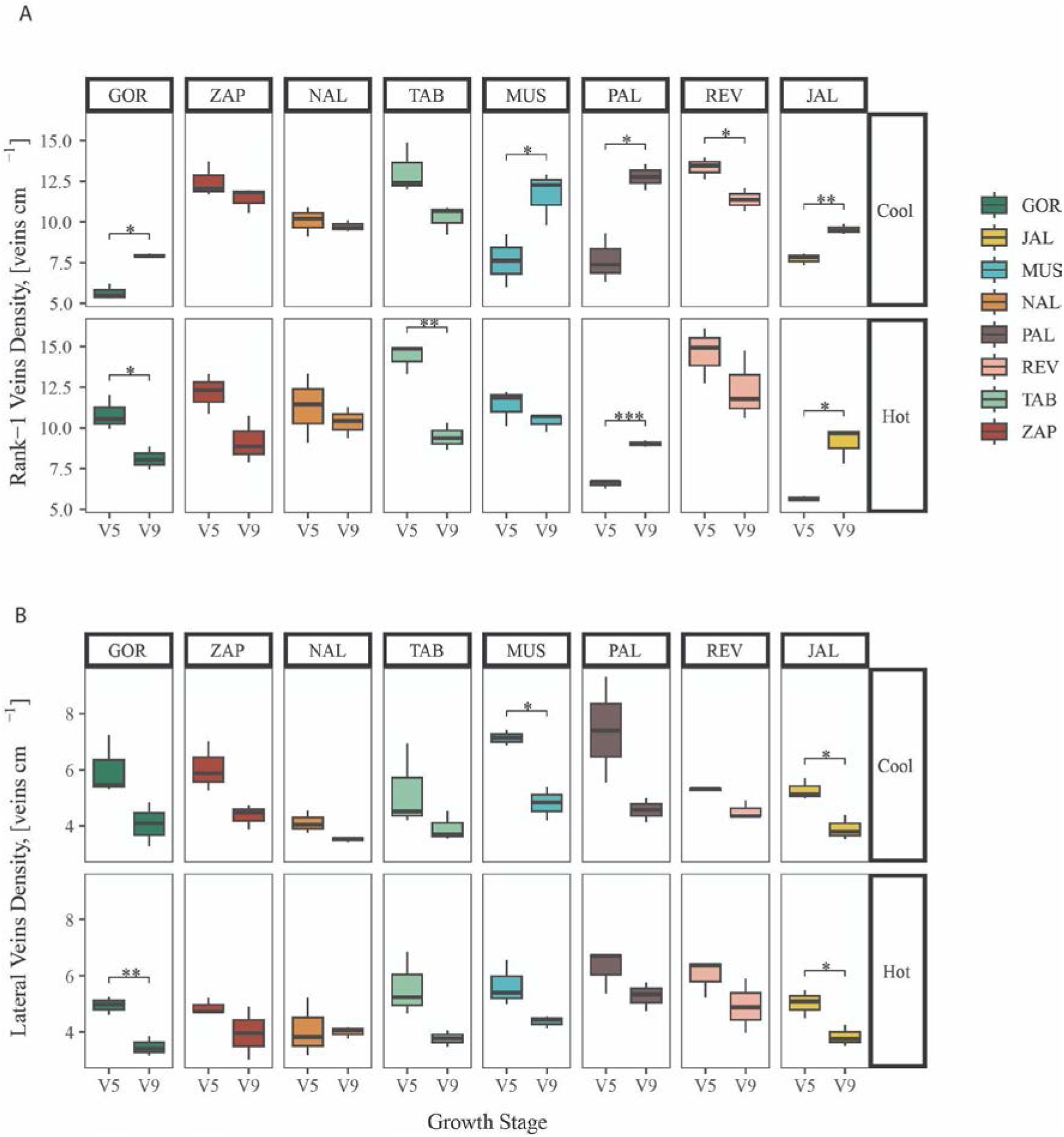
r1 and lateral vein density quantification across development and temperature. (A) r1 vein density at developmental stages V5 and V9 in two environmental conditions (Cool, 20°C; Hot, 30°C). (B) lateral vein density at developmental stages V5 and V9 in two environmental conditions (Cool, 20°C; Hot, 30°C).

**Fig. S3.**
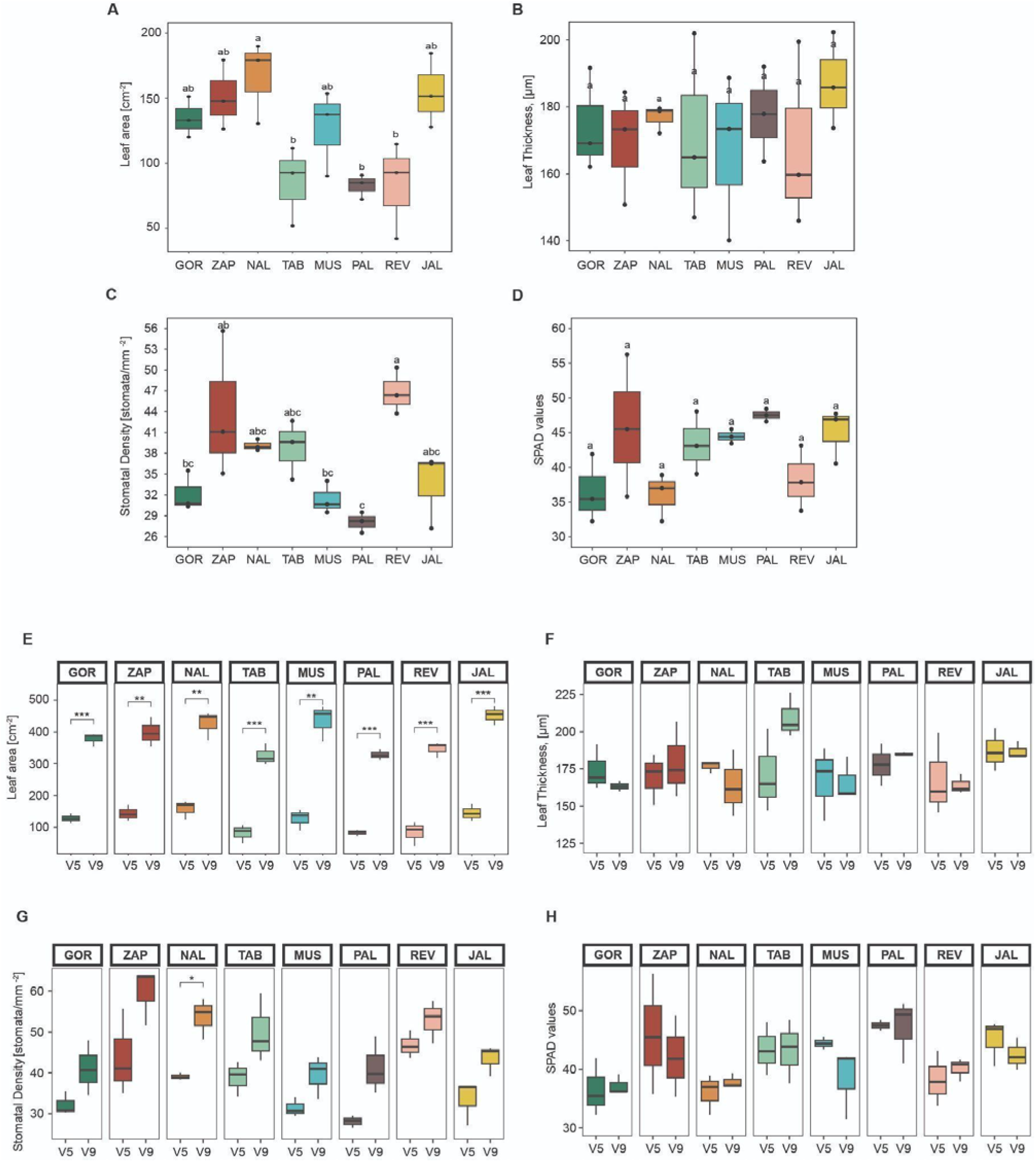
Morphological and relative chlorophyll content variation across genotypes. (A-D) Quantification of leaf morphological traits and chlorophyll relative content from plants grown in warm conditions. (E-F) Quantification of leaf traits across development from stage V5 to V9.

**Fig. S4.**
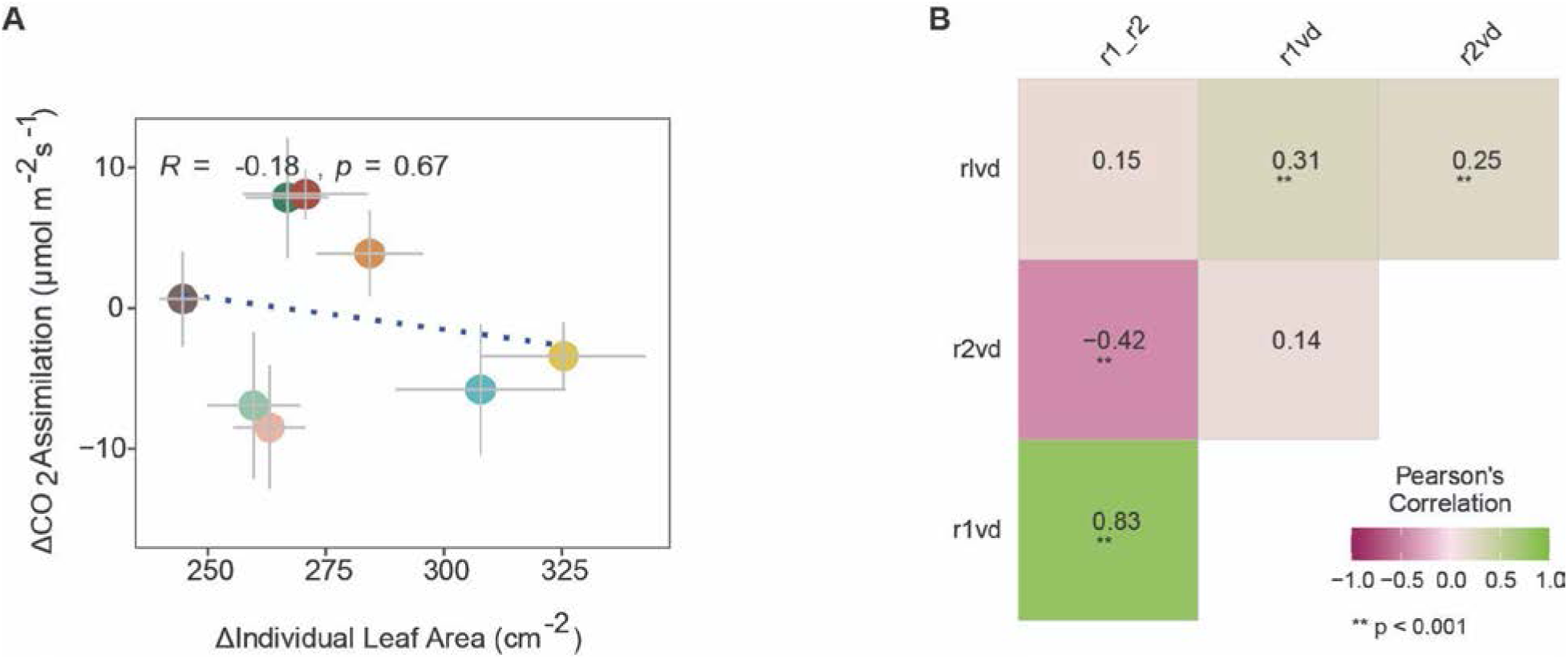
Correlation between phenotypic traits. (A) Pearson correlation and linear regression fit between the change in leaf area and CO_2_ assimilation values from V5 to V9 under heat. (B) Pearson correlation analysis between vein density trait values measured in the MAGIC population (BLUPs).

## Supplemental tables

**Table S1.** Photosynthesis and vein traits variables for stepwise linear regression.

**Table S2.** Hereditability estimates for vein density phenotypes.

**Table S3.** QTL coordinates and genes overlapping with each associated locus

**Table S4.** P4 gene expression normalized values and correlation coefficients for genes overlapping with r2vd QTLs.

**Table S5.** Accession information for the NMVs used in this study.

## Materials and Methods

### Plant material

Eight NMV were selected from distinct genotypic groups and geographic origins, based on its performance and contrasting environmental adaptation (Perez-Limón et al., 2022). Seeds were requested to and provided by the International Maize and Wheat Improvement Center (CIMMYT (https://www.genesys-pgr.org/wiews/MEX002). Detailed information about NMVs can be found in Table S5.

### Environmental variables at the NMVs geographic origins

We extracted data from minimum (Tmin) and maximum temperature (Tmax), precipitation (mm) (PREC), Potential Evapotranspiration (PET), Vapour Pressure (VP) and Vapour Pressure Deficit (VPD) from TERRACLIMATE, a global database with monthly average updates and a ∼4 km (1/24^th^ degree) spatial resolution (Abatzoglou et al., 2018) (https://www.climatologylab.org/terraclimate.html), covering the period from 1970-2020, and clipped-off to Mexico. We used *raster, terra, dplyr*, *purrr* and *sf* R packages to read raster files and convert them to dataframes. To get country outlines (Mexico) on the raster image we used *rnaturalearth* package (http://www.naturalearthdata.com). To retrieve the data points from each coordinate on the map (Figure 1A, 1C) we used AOI (https://github.com/mikejohnson51/AOI) and ClimateR (https://github.com/mikejohnson51/climateR) packages.

Mean Temperature (°C) (MT) and Mean Precipitation (mm) (MP) at the geographical origin of each NMV were calculated by averaging monthly values. The Aridity Index (AI) global raster data was retrieved from Global Aridity Index and Potential Evapotranspiration (ET0) Climate Database v3 (https://doi.org/10.6084/m9.figshare.7504448.v4) (Zomer et al., 2022) at 30 arc-seconds resolution (∼ 1km at the equator). Low values of AI indicates a drier climate. However we normalized AI values to perform PCA analyses such that PCA vectors showed a logical relationship with aridity, as described in the following equation:

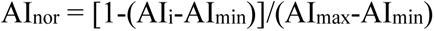

### Principal Components Analysis

Principal Components Analysis (PCA) was carried out to visualize the relationships among environmental variables at each NMV’s geographical origin (Aridity Index AI, Potential Evapotranspiration PET, Elevation Elev, Mean Temperature MT and Accumulated Precipitation AP) for the crop season period from May to October. PCA was performed using the FactoExtra package in R. The data were centred and scaled to account for differences in magnitude of the parameters.

### Plant growth conditions

An initial phenotyping screening of NMVs was done to determine if variation in photosynthetic traits existed by growing plants at the greenhouse of Unidad de Genómica Avanzada (UGA), Cinvestav, Irapuato, Mexico (N20°43′11.95′′, W101°19′50.78′′, 1730 masl). This greenhouse had an average temperature of 30 °C and 52 % relative humidity (RH) during the growing season. Seeds were first planted on 15 cm diameter pots with a mixture of peat moss and organic soil. Three replicates per landrace, and one seed per pot was germinated. Pots were irrigated every 3-4 days or as needed to keep the soil moist. Plants were then transferred to 50 cm diameter pots (30 days after germination) containing the same soil supplemented with Triple 17 Plus®. Fertilization was done once per month until plants reached maturity and tissue collection was done.

A second phenotyping experiment was done to determine plasticity in vein density traits in response to temperature. NMVs were grown in parallel under two conditions: a warm greenhouse with an average temperature of 28.1±1.0 °C and 56.4% ± 4.1 RH; and a cool greenhouse with an average temperature of 20.1±0.7 °C and 70.6% ± 4.5 RH. Seeds were planted by triplicate in the second week of June, 2022, in 20 liter free-draining pots containing a mixture of 3:1:1:4 peat moss, vermiculite, perlite and organic soil, respectively. Three replicates per landrace and two seeds per pot were germinated. Pots were watered as needed to keep the soil moist, and fertilized once every month with Triple 17 Fertilizer (Vigoro® Excelso). Plants were grown on a 13/11 hours day/night cycle (+23 to -44 min in day length along the growth cycle), for a total of 80 days for warm conditions and 96 for cold conditions. Plants in both conditions received the approximately same total growing degree days (approx. 1900 GDD).

### Leaf tissue sampling

For NMVs, we collected the middle portion of fully expanded and healthy leaves at different developmental stages: leaf 5 at V5 and leaf 9 at V9, corresponding to 31 and 62 days after planting, respectively. Three biological replicates (leaves from different plants) from each NMV were sampled, labeled and stored at 4 °C in 70% ethanol until processed.

In the case of the MexMAGIC population sampled at the field, we took the middle portion of the 3rd leaf below the tassel. Two replicates (plants) per each row (family) at the tasseling stage (VT) were included. Once cut, the tissue was rolled and stored in labeled 50 ml Falcon tubes containing a solution of ethanol: distillated water (70% v/v). Then the samples were transported to the laboratory and stored at 4°C until processed.

### Histological analysis of leaf sections

Leaf tissue stored in ethanol at 4°C was manually dissected into smaller sections of approximately 10 mm large and ∼0.5 mm wide with a scalpel, encompassing the mid-vein and 4-5 adjacent lateral veins. Tissue was then stained in a solution of 0.3 mg/L^-1^ Calcofluor White Stain (Sigma-Aldrich, cat. no. 18909) for 60 seconds (Atkinson and Wells, 2017). After being rinsed in distillate water, samples were positioned upright (veins vertically oriented) into 1 cm^2^ molds and filled with 7% agar (Sigma-Aldrich, cat. no. A1296). Once cooled, blocks were trimmed (∼1-2 mm) using a cutter and mounted such that veins were vertically orientated. Blocks were placed on a microscope slide with distilled water and covered with a coverslip (see Fig. S1)

### Image acquisition and processing

Leaf sections were imaged immediately, using a Leica DM6000B fluorescence microscope with Leica DMC6200 camera (Leica Microsystems CMS GmbH, https://www.leica-microsystems.com) and Leica Application Suite v4.13.0 [Build:167] software (Leica Microsystems, https://www.leica-microsystems.com), and saved in TIFF or JPG format. Images were taken using both the “A” (UV) and “I3” (blue) filter cubes (Excitation filter: BP340-380 and BP 450-490, respectively), which enabled visualization of cell walls stained with calcofluor and sclerenchyma respectively. Analyses of both images facilitated vein type identification and quantification. Images were acquired at a total magnification of 50x.

Because leaf sections generated by manually cutting with a scalpel were not completely even, a section of the image in the field of view often appeared out of focus. To solve this issue, we took several photographs maintaining the field of view but changing depth to achieve optimal focus in all image areas. We then used the *PMax* image stacking technique by Zerene Stacker software v1.04 (64bit) (Zerene Systems LLC, https://www.zerenesystems.com/), to stack several images combining them into a single focused image. To accelerate the process of stacking, a Python™ script was developed. In addition, several images per sample were acquired to compose one large micrograph of a single leaf. Image stitching was done by overlapping the focus-corrected images to produce a panoramic image using Autostich (http://matthewalunbrown.com/autostitch/autostitch.html).

Subsequent quantification of vein traits was done using the ImageJ software package (National Institutes of Health, https://imagej.nih.gov/). A macro was used to automatically obtain the name of the file, set the scale in µm and save the results datasheet in a csv file. To accelerate the process of distance measurement we used a VINSA 1060 Plus Graphics Drawing Tablet 10*6 inches (http://www.ping-it.cn/).

### Quantification of leaf anatomical traits

Leaf veins were classified as belonging to three types: lateral veins (lv), rank-1 veins (r1) and rank-2 veins (r2). Lateral veins were distinguished by their size and vasculature diameter, which was notably bigger compared to r1 and r2 veins. r1 veins were distinguished by the presence of sclerenchyma and the space occupied between upper and lower epidermis (Sedelnikova et al., 2018), and r2 veins were similar in size compared to r1 but without sclerenchyma. One section of each leaf image from the midrib to the margin was measured and vein types quantified to get estimates of vein density.

Leaf Thickness was assessed using the same photos that were used for vein density quantification. We averaged three measurements for each individual photo, and three photos along the medio-lateral axis to include possible variation across areas near and far to the mid-vein for each replicate (plant) and all the varieties.

Stomatal number was determined from images obtained from imprints of abaxial and adaxial surfaces of leaves. Nail varnish was used to produce impressions of the leaf epidermis. Once dry, the varnish was peeled off by using clear tape and immediately pressed onto a microscope slide. For quantification we considered a FOV of 2250*1406 μm = 3.1635 mm^2^ at 50X magnification and used this area to calculate stomatal density (mm^-2^). Values are averages of three biological replicates per NMV. Since the stomatal frequency decreases from the tip to the base of the leaf surface of maize (Heichel, 1971), measurements were done next to the midpoint (the same area where tissue for histological analysis was taken) of the proximal-distal axis of the leaf.

### Photosynthesis measurements and Response Curves

Steady state maximum photosynthetic rates (A_max_) and A/Ci curves were generated using an Infra-Red Gas Analyzer (IRGa) (CIRAS-3, PP Systems, Inc., Amesbury, MA, USA) with a 32 cm^2^ PLC3 leaf cuvette and a PLC3 LED Light Unit (RGBW) to control the illumination.

Amax measurements were made on fully expanded L5 between 30 and 40 DAP, and for L9 between 50 and 60 DAP at a photosynthetic photon flux density (PPFD) of 1000 *μ*mol photons m^-2^ s^-1^, 40% humidity and 30°C or 20 °C leaf temperature (set at the leaf cuvette). In all cases the flow rate was set in 300 *μ*mol s^-1^. Leaves were clamped at the midpoint along the proximal-distal axis and allowed to acclimate for at least 5 minutes at 400 *μ*mol mol^-1^ CO_2_ until photosynthetic rate had stabilized.

A/Ci curves were generated by measuring photosynthetic rates at 400, 300, 200, 100, 50, 25, 400 (recovery, omitted from curve plotting), 600, 800, and 1000 *μ*mol mol^-1^ CO_2_. At each new CO_2_ concentration, measurements were only taken at steady state, normally between 2 and 6 minutes after the desired CO_2_ concentration was achieved. All measurements were recorded three times (10 seconds between records) and averaged. Measurements were taken throughout the day between 11:00 am and 16:00 pm, with genotypes randomly sampled to avoid any circadian influences. Leaf area within the cuvette was adjusted, measuring the width of leaf and multiplying by the length of cuvette. Individual A/Ci curves were modeled using the equations described previously for C_4_ photosynthesis (von Caemmerer, 2000). The most recent kinetic constants described for maize were used in order to fit the model(Zhou et al., 2019), with PhotoGEA v0.10.0/R, (https://eloch216.github.io/PhotoGEA/index.html), and to analyze C_4_ A/Ci curves, combined with C4-Parameter-Estimation (https://rdrr.io/github/zhouhaoran06/C4-Parameter-Estimation/) (Zhou et al., 2019). Modeled data agreed with experimental data.

### Chlorophyll relative content

Measurements of chlorophyll relative content were made using a SPAD chlorophyll meter (SPAD 502 Plus, Konica Minolta Inc., Tokyo, Japan. (https://www.konicaminolta.com/instruments/download/catalog/color/pdf/spad502plus_catalog_eng.pdf) on fully expanded leaves 5 and 9. Readings were taken at the center and tip of the leaf adaxial side avoiding the midribs. Three readings were taken for each leaf position, and an average value per leaf was reported.

### Stepwise linear regression model

To find the best predictor variables for a linear model in which maximum photosynthetic rates were the dependent variables, the StepReg package was used (https://CRAN.r-project.org/package=StepReg). A bidirectional linear stepwise regression model with the AIC metric was applied including the following variables: dAmax ∼ R2VD_V5 + R2VD_V9 + dR2VD + R1VD_V5 + R1VD_V9 + RLVD_V5 + RLVD_V9 + dR1VD + dR1_R2 + dRLVD + R1_R2_V5 + R1_R2_V9 (Table S4). Best predictor was determined based on coefficients significance.

### Generation of MexMAGIC families

A maize MexMAGIC population (Yu et al., 2024b, Yu et al., 2024a) was generated by crossing eight selected founders (5 outbred varieties; 3 S_6_ inbred lines) following a diallel scheme and a set of four successful F_1_ combinations selected to cover all eight parents. Subsequently, four-way crosses were made using a single individual from each F_1_ to capture a single haplotype per founder using the “chain” scheme A to B, B to C, C to D and D to A. An equal number of seeds was advanced from each four-way ear for a round of eight-way crosses using multiple individuals, followed by 3 generations of random intermating, and 3 generations of inbreeding. For simplicity, the final families were designated Intermated3 Selfed3 (I_3_S_3_). The I_3_S_3_ MexMAGIC were test-crossed to the inbred line NC358 for following field evaluations. More details of population generation are given in Supplemental Material.

### Parental assignment across the genomes of MexMAGIC families

To characterize the genetic mosaic of the MexMAGIC families and to saturate the physical map, a subset of 1M random SNPs was selected with stratification by chromosome using the *R/rsample::initial_split* function (https://rsample.tidymodels.org/reference/initial_split.html). To provide a genetic position to the SNPs in the MexMAGIC dataset, the package *R/scam* (Pya and Wood, 2015) was used to train a monotonic smooth with the physical and genetic position of the SNPs in the IBM RIL maize population (Lee et al. 2002), and use it to predict the genetic position on the MexMAGIC dataset. The genetic and physical map, and the genotypes of both the founder and MexMAGIC families were formatted into a cross object to be used by the *R/qtl2* package (Broman et al., 2019). The founder conditional genotype probabilities at every marker were estimated with the *qtl2::calc_genoprob* function and the founder genotype with maximum marginal probability was estimated with the *qtl2::maxmarg*.

### QTL and SNP-association mapping, estimation of allelic effect

To map functional variation in leaf anatomic traits, segregation in the MexMAGIC founders, a SNP-Association (GWAS) and haplotype-based QTL mapping methods available in *R/qtl2* (Broman et al., 2019) were used to estimate the genetic relation between families and to control for residual population structure in the models. A “leave one chromosome out” (LOCO) kinship matrix was estimated using *qtl2::calc_kinship()*. The previously estimated founder conditional genotype probabilities were transformed into allele probabilities using *qtl2::genoprob_to_alleleprob()* function. To estimate the association between phenotypes and biallelic SNPs segregating in the founders of the MexMAGIC population, a linear mixed model was fitted using the *qtl2::scan1snps()* function, using as inputs the allele probabilities, phenotypic information, and the kinship matrix to account for residual population structure. The results of the SNP-association were summarised in Manhattan plots using *R/ggplot2* package for R (Wickham, 2009). To understand the effect of the association between the segregation of phenotypes and the effect of the founder alleles, a single-QTL scan was performed using an 8-allele method. A linear mixed model was fitted using the *qtl2::scan1()* function, using as inputs the marginal genotype probabilities, phenotypic information and the kinship matrix to account for residual structure. The LOD trace plots were visualized using the *R/ggplot2* package. To establish a threshold to detect significant signals, a 1000 permutation test was run using the function *qtl2::scan1perm()* function considering a significance threshold (alpha) of 0.1. Significant QTLs were identified with the function *qtl2::find_peaks()* and the LOD-support interval was considered as a drop of 2 in LOD value from the QTL peak (Broman et al., 2019). To obtain an estimation and confidence interval of the allelic founder effect at every QTL, the marginal genotype probabilities at the lead SNP were extracted using *ql2::pull_genoprobpos()* function. The subsetted genotype probabilities and phenotypic data were used as input for the *qtl2::fit1()* function to fit a single-QTL model using a mixed linear model and kinship matrix to control for residual structure. The estimated coefficient and standard errors of the founder allelic effect were extracted from the QTL object and transformed into a tibble for posterior analysis and visualization.

### Field phenotypic evaluation of MexMAGIC families

The MexMAGIC population plus three test controls (B73, Mo17, and CML312 inbred lines) were planted in a complete block of 240 families at Valle de Banderas, Nayarit, México (N20°47′41.37′′, W105°14′51.49′′, 51 masl) during the winter of 2020/2021. The distribution of the families was randomized. 22 plants per family were sown in 2 m long rows. Plants were grown under standard agronomic conditions with irrigation.

### RNA isolation and sequencing

The NMVs were grown in parallel in two greenhouses under an average temperature of 29 °C and 20 °C and relative humidity ranging from 44.5% to 64%. The fifth (V5) and ninth (V9) leaves were collected at developmental stage P4 (plastochron 4). Total mRNA was extracted using the RNeasy Mini Kit (QIAGEN), following the manufacturer’s instructions, including on-column DNA digestion step (Turbo DNase, Thermo Fisher). Two biological replicates were used, and 4 to 5 leaf primordia were harvested for each replicate. Library preparation was performed using the NEBNext Ultra II Directional RNA Library Prep Kit for Illumina. A total of 32 libraries were sequenced on the Illumina NovaSeq 6000 platform, generating 150 bp paired-end reads. Read alignment was performed using STAR (version 2.7.3a) against the Zm-B73-REFERENCE-NAM-5.0 reference genome. Mapped read counts were obtained using FeatureCounts.

### Transcriptome to phenotype correlation analyses

Expression values were normalized using DESeq2 with default parameters. A total of 32 samples were analyzed, corresponding to two developmental stages and eighth NMVs. To obtain the expression deltas, the following procedure was applied: the difference in expression between stage V5 and stage V9 was calculated individually for each replicate, these values were then averaged to generate an average delta. The same procedure was applied to obtain vein density deltas. Averaged deltas were then used to calculate correlation analyses using the Pearson correlation coefficient. Gene selection was based on two criteria: (i) genes with a p-value ≤ 0.05 and a positive Pearson correlation coefficient > 0.5, and (ii) genes with a p-value ≤ 0.05 and a negative Pearson correlation coefficient < –0.5. For the construction of the correlogram, the top 10 genes showing the strongest positive and negative correlations were considered, with Pearson coefficient above or below 0.7. We calculated the founder allelic effects for r2vd at the leading SNPs for each of the 20 top genes and calculated Pearson correlation coefficients between the parental effects and environmental variables (Tmean, VP, VPD) and gene expression values. Genes with significant association were highlighted.

## Author contributions

JLCR, COR and RJHS designed the study. JLCR conducted the greenhouse experimental work and generated the phenotype and histological data. RJHS provided the MexMAGIC population. ANAN and RJHS conducted field work. JLCR, COR, SPL and ANAN collected leaf tissue at the field. JLCR collected leaf tissue and performed photosynthesis measurements. DCE helped with obtaining cross sections and analyzing micrographs. JLCR analyzed phenotypic and histological data. DCE helped with the analysis and interpretation of photosynthetic parameters. JLCR and SPL performed mapping of genetic regions. EHJ collected plastochron 4 tissue, extracted RNA, prepared libraries and analyzed gene expression data. NGC helped with the analyses of candidate genes. JLCR, COR and MHC interpreted results and wrote the manuscript, and MJOE assisted in greenhouse experiments.

## Acknowledgments

JCR and DCE are supported by a PhD scholarship from CONACYT-Mexico; NGC is supported by a MSc scholarship by CONACYT.

## Data availability

Sequencing data supporting the conclusions of this paper has been deposited in the Gene Expression Omnibus (GEO) under accession GSE314465.

## References

Abatzoglou, J. T., Dobrowski, S. Z., Parks, S. A. & Hegewisch, K. C. 2018. TerraClimate, a high-resolution global dataset of monthly climate and climatic water balance from 1958-2015. Sci Data, 5, 170191.

Alenazi, A. S., Pereira, L., Christin, P. A., Osborne, C. P. & Dunning, L. T. 2024. Identifying genomic regions associated with C(4) photosynthetic activity and leaf anatomy in Alloteropsis semialata. New Phytol, 243, 1698–1710.

Amiard, V., Mueh, K. E., Demmig-Adams, B., Ebbert, V., Turgeon, R. & Adams, W. W., 3rd 2005. Anatomical and photosynthetic acclimation to the light environment in species with differing mechanisms of phloem loading. Proc Natl Acad Sci U S A, 102, 12968–73.

Atkinson, J. A. & Wells, D. M. 2017. An Updated Protocol for High Throughput Plant Tissue Sectioning. Front Plant Sci, 8, 1721.

Boyce, C. K., Brodribb, T. J., Feild, T. S. & Zwieniecki, M. A. 2009. Angiosperm leaf vein evolution was physiologically and environmentally transformative. Proc Biol Sci, 276, 1771–6.

Brodribb, T. J., Feild, T. S. & Jordan, G. J. 2007. Leaf maximum photosynthetic rate and venation are linked by hydraulics. Plant Physiol, 144, 1890–8.

Broman, K. W., Gatti, D. M., Simecek, P., Furlotte, N. A., Prins, P., Sen, S., Yandell, B. S. & Churchill, G. A. 2019. R/qtl2: Software for Mapping Quantitative Trait Loci with High-Dimensional Data and Multiparent Populations. Genetics, 211, 495–502.

Cohu, C. M., Muller, O., Stewart, J. J., Demmig-Adams, B. & Adams, W. W. 2013. Association between minor loading vein architecture and light- and CO-saturated rates of photosynthetic oxygen evolution among ecotypes from different latitudes. Frontiers in Plant Science, 4.

Dhakal, S., Reiter, J. W., Laroche, A. & Schultz, E. A. 2021. Leaf vein pattern response to heat and drought requires genes that influence PINFORMED1 localization and is mimicked by ABA treatment. Environmental and Experimental Botany, 185.

Feldman, A. B., Leung, H., Baraoidan, M., Elmido-Mabilangan, A., Canicosa, I., Quick, W. P., Sheehy, J. & Murchie, E. H. 2017. Increasing Leaf Vein Density via Mutagenesis in Rice Results in an Enhanced Rate of Photosynthesis, Smaller Cell Sizes and Can Reduce Interveinal Mesophyll Cell Number. Front Plant Sci, 8, 1883.

Garthwaite, I. J., Lepp, C., Maldonado, Z. S. R., Blasini, D., Grady, K. C., Gehring, C. A., Hultine, K. R., Whitham, T. G., Allan, G. J. & Best, R. J. 2024. Plasticity in Hydraulic Architecture: Riparian Trees Respond to Increased Temperatures With Genotype-Specific Adjustments to Leaf Traits. Ecol Evol, 14, e70683.

Gowik, U. & Westhoff, P. 2011. The path from C3 to C4 photosynthesis. Plant Physiol, 155, 56–63.

Harrison, E. L., Arce Cubas, L., Gray, J. E. & Hepworth, C. 2020. The influence of stomatal morphology and distribution on photosynthetic gas exchange. Plant J, 101, 768–779.

Heichel, G. H. 1971. Genetic Control of Epidermal Cell and Stomatal Frequency in Maize. Crop Science, 11, 830–&.

Hu, W., Lu, Z., Meng, F., Li, X., Cong, R., Ren, T., Sharkey, T. D. & Lu, J. 2020. The reduction in leaf area precedes that in photosynthesis under potassium deficiency: the importance of leaf anatomy. New Phytol, 227, 1749–1763.

Hughes, T. E., Sedelnikova, O., Thomas, M. & Langdale, J. A. 2023. Mutations in NAKED-ENDOSPERM IDD genes reveal functional interactions with SCARECROW during leaf patterning in C4 grasses. PLoS Genet, 19, e1010715.

Hughes, T. E., Sedelnikova, O. V., Wu, H., Becraft, P. W. & Langdale, J. A. 2019. Redundant SCARECROW genes pattern distinct cell layers in roots and leaves of maize. Development, 146.

Jiang, A., Guo, Z., Pan, J., Yang, Y., Zhuang, Y., Zuo, D., Hao, C., Gao, Z., Xin, P., Chu, J., Zhong, S. & Li, L. 2021. The PIF1-miR408-PLANTACYANIN repression cascade regulates light-dependent seed germination. Plant Cell, 33, 1506–1529.

Karki, S., Rizal, G. & Quick, W. P. 2013. Improvement of photosynthesis in rice (Oryza sativa L.) by inserting the C4 pathway. Rice (N Y), 6, 28.

Liu, Q., Teng, S., Deng, C., Wu, S., Li, H., Wang, Y., Wu, J., Cui, X., Zhang, Z., Quick, W. P., Brutnell, T. P., Sun, X. & Lu, T. 2023. SHORT ROOT and INDETERMINATE DOMAIN family members govern PIN-FORMED expression to regulate minor vein differentiation in rice. Plant Cell, 35, 2848–2870.

Liu, W. Y., Chang, Y. M., Chen, S. C., Lu, C. H., Wu, Y. H., Lu, M. Y., Chen, D. R., Shih, A. C., Sheue, C. R., Huang, H. C., Yu, C. P., Lin, H. H., Shiu, S. H., Ku, M. S. & Li, W. H. 2013. Anatomical and transcriptional dynamics of maize embryonic leaves during seed germination. Proc Natl Acad Sci U S A, 110, 3979–84.

Lundgren, M. R., Dunning, L. T., Olofsson, J. K., Moreno-Villena, J. J., Bouvier, J. W., Sage, T. L., Khoshravesh, R., Sultmanis, S., Stata, M., Ripley, B. S., Vorontsova, M. S., Besnard, G., Adams, C., Cuff, N., Mapaura, A., Bianconi, M. E., Long, C. M., Christin, P. A. & Osborne, C. P. 2019. C4 anatomy can evolve via a single developmental change. Ecol Lett, 22, 302–312.

Osborne, C. P. & Sack, L. 2012. Evolution of C4 plants: a new hypothesis for an interaction of CO2 and water relations mediated by plant hydraulics. Philos Trans R Soc Lond B Biol Sci, 367, 583–600.

Perico, C., Zaidem, M., Sedelnikova, O., Bhattacharya, S., Korfhage, C. & Langdale, J. A. 2024. Multiplexed in situ hybridization reveals distinct lineage identities for major and minor vein initiation during maize leaf development. Proc Natl Acad Sci U S A, 121, e2402514121.

Pya, N. & Wood, S. N. 2015. Shape constrained additive models. Statistics and Computing, 25, 543–559.

Rishmawi, L., Buhler, J., Jaegle, B., Hulskamp, M. & Koornneef, M. 2017. Quantitative trait loci controlling leaf venation in Arabidopsis. Plant Cell Environ, 40, 1429–1441.

Robil, J. M. & Mcsteen, P. 2023. Hormonal control of medial-lateral growth and vein formation in the maize leaf. New Phytol, 238, 125–141.

Sedelnikova, O. V., Hughes, T. E. & Langdale, J. A. 2018. Understanding the Genetic Basis of C4 Kranz Anatomy with a View to Engineering C3 Crops. Annu Rev Genet, 52, 249–270.

Simpson, D. E. O. S., , G. R., M. Eric Schranz , & Pallavi Singh, J. M. H. 2025. Genetic Mapping for Agronomic, Nutritional, and Leaf Vein Traits in the Indigenous Crop 2 Gynandropsis gynandra. Sustainable agriculture, 3.

Slewinski, T. L., Anderson, A. A., Price, S., Withee, J. R., Gallagher, K. & Turgeon, R. 2014. Short-root1 plays a role in the development of vascular tissue and kranz anatomy in maize leaves. Mol Plant, 7, 1388–1392.

Vlad, D., Zaidem, M., Perico, C., Sedelnikova, O., Bhattacharya, S. & Langdale, J. A. 2024. The WIP6 transcription factor TOO MANY LATERALS specifies vein type in C(4) and C(3) grass leaves. Curr Biol, 34, 1670–1686 e10.

Von Caemmerer, S. 2000. Biochemical Models of Leaf Photosynthesis, Collingwood:, CSIRO Pub.

Von Arx, G., Archer, S. R. & Hughes, M. K. 2012. Long-term functional plasticity in plant hydraulic architecture in response to supplemental moisture. Ann Bot, 109, 1091–100.

Wang, J., Lu, W., Tong, Y. & Yang, Q. 2016. Leaf Morphology, Photosynthetic Performance, Chlorophyll Fluorescence, Stomatal Development of Lettuce (Lactuca sativa L.) Exposed to Different Ratios of Red Light to Blue Light. Front Plant Sci, 7, 250.

Wang, X., Chen, S., Yang, X., Zhu, R., Liu, M., Wang, R. & He, N. 2024. Adaptation mechanisms of leaf vein traits to drought in grassland plants. Sci Total Environ, 917, 170224.

Wickham, H. 2009. ggplot2: Elegant Graphics for Data Analysis. Ggplot2: Elegant Graphics for Data Analysis, 1–212.

Yang, Y., Xu, L., Hao, C., Wan, M., Tao, Y., Zhuang, Y., Su, Y. & Li, L. 2024. The microRNA408-plantacyanin module balances plant growth and drought resistance by regulating reactive oxygen species homeostasis in guard cells. Plant Cell, 36, 4338–4355.

Ye, M., Wu, M., Zhang, H., Zhang, Z. L. & Zhang, Z. J. 2021. High Leaf Vein Density Promotes Leaf Gas Exchange by Enhancing Leaf Hydraulic Conductance in L. Plants. Frontiers in Plant Science, 12.

Yu, B., Zhou, C., Wang, Z., Bucher, M., Schaaf, G., Sawers, R. J. H., Chen, X., Hochholdinger, F., Zou, C. & Yu, P. 2024a. Maize zinc uptake is influenced by arbuscular mycorrhizal symbiosis under various soil phosphorus availabilities. New Phytol, 243, 1936–1950.

Yu, P., Li, C., Li, M., He, X., Wang, D., Li, H., Marcon, C., Li, Y., Perez-Limon, S., Chen, X., Delgado-Baquerizo, M., Koller, R., Metzner, R., Van Dusschoten, D., Pflugfelder, D., Borisjuk, L., Plutenko, I., Mahon, A., Resende, M. F. R., Jr., Salvi, S., Akale, A., Abdalla, M., Ahmed, M. A., Bauer, F. M., Schnepf, A., Lobet, G., Heymans, A., Suresh, K., Schreiber, L., Mclaughlin, C. M., Li, C., Mayer, M., Schon, C. C., Bernau, V., Von Wiren, N., Sawers, R. J. H., Wang, T. & Hochholdinger, F. 2024b. Seedling root system adaptation to water availability during maize domestication and global expansion. Nat Genet, 56, 1245–1256.

Zhou, H., Akcay, E. & Helliker, B. R. 2019. Estimating C(4) photosynthesis parameters by fitting intensive A/C(i) curves. Photosynth Res, 141, 181–194.

Zomer, R. J., Xu, J. & Trabucco, A. 2022. Version 3 of the Global Aridity Index and Potential Evapotranspiration Database. Sci Data, 9, 409.

